# Remodeling of RNA-Binding Proteome and RNA-mediated regulation as a new layer of control of sporulation

**DOI:** 10.1101/2025.03.21.644579

**Authors:** Thomas Kaboré, Maya Belghazi, Christophe Verthuy, Anne Galinier, Clémentine Delan-Forino

## Abstract

Sporulation allows certain bacteria to survive extreme conditions for extended periods posing challenges to public health and food safety. Transcriptional level of regulation relying on σ factors has been well studied in the non-pathogenic model bacterium *Bacillus subtilis*, while post-transcriptional control remains poorly understood. RNA-binding proteins (RBPs) and small non-coding RNAs (sRNAs) modulate gene expression by affecting mRNA stability or translation. Recent studies suggest that RNA-mediated regulation plays a role in sporulation, but its networks and interactions remain largely uncharacterized, highlighting the need for further investigation. To address this knowledge gap, we adapted OOPS (Orthogonal Organic Phase Separation), a high-throughput method to specifically purify RNA binding proteome (RBPome). By monitoring its dynamics, we observed RBPome is highly remodeled during *B. subtilis* process. Our approach led to the identification of novel RBPs and potential RNA-mediated post-transcriptional regulators of sporulation. This work provides important insights into the interplay between RNAs and RBPs, advancing the understanding of post-transcriptional regulation in Gram-positive spore-forming bacteria. It also offers new resources for exploring the molecular mechanisms that govern sporulation.

**IMPORTANCE:** Understanding how bacteria survive extreme conditions is key to tackling challenges in health, food safety, and industry. This study reveals a previously unexplored layer of control in *Bacillus subtilis*, a model organism for spore-forming bacteria—those that can produce spores, which are highly resistant, dormant cells able to endure harsh environments. By mapping, for the first time, the full set of proteins that interact with RNA during sporulation, this work uncovers how bacteria fine-tune their internal programs. The identification of novel RNA-binding proteins sheds light on how bacteria adapt at the molecular level and lays a valuable foundation for future mechanistic studies that will deepen our understanding of bacterial adaptation and resilience.

## INTRODUCTION

Bacteria can adapt to changing environments and survive extreme conditions. For example, under nutrient-limiting conditions, members of the Bacillus and Clostridium genera can undergo sporulation, differentiating into highly resilient dormant spores that enter extreme quiescence (Stragier 2022). When conditions improve, spores germinate and revert to metabolically active cells, resuming vegetative growth even after 250 million years in some cases (Vreeland et al. 2000). This makes them some of the longest-living cellular forms.

*Bacillus subtilis* is a rod-shaped bacterium, ubiquitously found in soil, and one of the best-characterized Gram-positive spore-forming bacteria (Higgins and Dworkin 2012). *Bacilli* sporulation is a well-defined multistage process (Fig. 1) (Riley et al. 2021). First, an asymmetric septum is formed, dividing the progenitor cell into a small forespore (the future spore) and a larger mother cell that supports spore maturation. Subsequently, the forespore is engulfed by the mother cell and a thick cell wall material (the cortex), is synthesized around it, followed by an inner and outer proteinaceous coat and a glycoprotein crust, shielding it from the extracellular surrounding. Dehydration of the spore core further contributes to cellular dormancy, a process unfolding over days while remaining responsive to extracellular signals. Mechanisms controlling protein synthesis and enzymatic activity during sporulation have been widely investigated in *B. subtilis*, particularly the role of σ factors in orchestrating gene transcription (Fig. 1) (Haldenwang and Losick 1979; Haldenwang et al. 1981; Kroos et al. 1989; Eichenberger et al. 2004; Hilbert and Piggot 2004). These alternative σ-factors lead the core RNA polymerase to different sets of promoters from those recognized by the housekeeping σ^A^ (Fig. 1); they control more than 500 genes, most of them are not being expressed during vegetative growth. Nearly a third of them remain functionally uncharacterized and even those with assigned function in spore formation are mostly partially understood (Zhu and Stülke 2018). To date, most of the researches have focused on the characterization of “*spo*” genes whose deletion affects sporulation without disturbing vegetative growth (Riley et al. 2021) and post-transcriptional regulation of sporulation remains largely unexplored. It usually relies on dynamic interactions between RNA-binding proteins (RBPs) and their RNA targets, known to regulate the adaptative response across all domains of life by modulating RNA metabolism, cellular homeostasis, and stress adaptation (Holmqvist and Vogel 2018; Delan-Forino et al. 2020; Trendel et al. 2019; Bresson et al. 2020; Goswami et al. 2024).

**Fig. 1:**
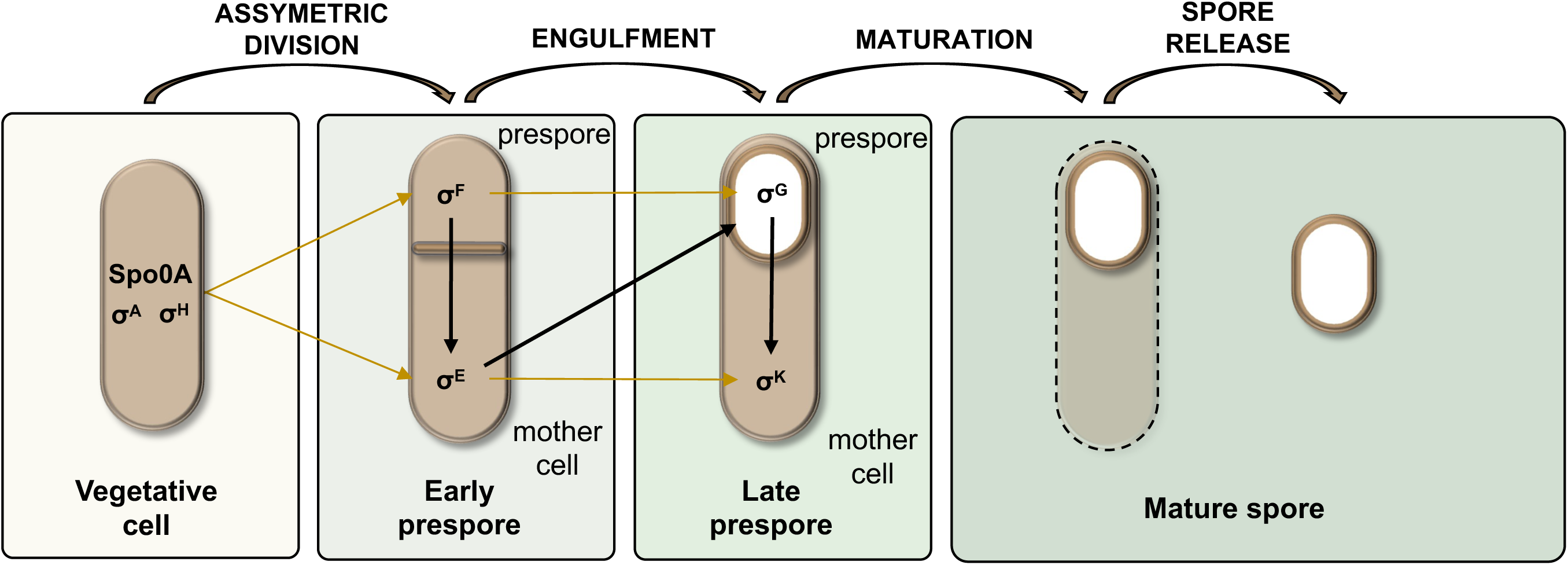
Overview of sporulation with cell-specific (pre-spore or mother cell) σ transcription factors involved in each step (serial activation represented as arrows). σ factors are subunits of the RNA polymerase responsible for determining the specificity of promoter DNA binding and required for transcription initiation.

In bacteria, RBPs often cooperate with regulatory RNAs such as small non-coding RNAs (sRNAs) to influence translation or transcript stability, mediating sRNA-mRNA interactions and/or recruit degradation factors (Christopoulou and Granneman 2022). sRNAs, transcribed transiently in response to environmental shifts (Nicolas et al. 2012), work with RBPs to target one or multiple mRNAs to fine-tune various cellular processes (Jørgensen et al. 2020) (Holmqvist and Wagner 2017). Recently, their involvement in sporulation has gained attention. For instance, SR1, a dual-function sRNA, targets *kinA* mRNA, encoding a kinase controlling sporulation initiation in *B. subtilis*. Other factors such as CsfG, conserved among spore-forming bacteria, or KrrA, a conserved RBP identified in *B. anthracis* (Pi et al. 2022; Marchais et al. 2011), have been implicated in regulation of mRNA involved in sporulation, germination and competence.

Unlike Gram-negative bacteria, where general RBPs such as Hfq and ProQ facilitate sRNA-mRNA interactions (Christopoulou and Granneman 2022), *B. subtilis* lacks ProQ, and Hfq appears dispensable for sRNA-mediated regulation (Panda et al. 2015; Yu et al. 2022; Rochat et al. 2015). Instead, Gram-positive bacteria likely rely on distinct, process-specific RBPs, such as CsrA, a pleiotropic regulator that stabilizes SR1 interactions with *ahrC* but not *kinA* (Müller et al. 2019).

Emerging methods such as poly-A-independent RBP purification have facilitated RBP discovery in prokaryotes. For example, Grad-seq, which combines density gradient ultracentrifugation with high-throughput sequencing, led to the identification of ProQ (Smirnov et al. 2016). While powerful, this technique does not confirm direct RNA–protein interaction. Historically, UV irradiation has been a robust tool to capture direct protein–RNA interactions by inducing covalent crosslinking. UV-crosslinking based methods, such as TRAPP (Shchepachev et al. 2019), 2C (Chu et al. 2022) leverage the intrinsic affinity of RNA for silica, while others, like PTex (Urdaneta et al. 2019) and Orthogonal Organic Phase Separation (OOPS) (Queiroz et al. 2019) rely on organic extraction. These approaches have successfully identified new RNA partners and regulatory pathways across diverse organisms (yeast, human cells, *E. coli*, *Staphylococcus aureus*) and can isolate crosslinked protein–RNA complexes, enabling the study of RBP dynamics.

Here, we adapted OOPS to *B. subtilis,* isolating the RNA-Binding proteome (RBPome) for the first time in a Gram-positive bacterium. We followed its dynamics during sporulation and demonstrated that both RBP production levels and RNA-binding abilities are highly modulated throughout this process. By this *in vivo* global approach, we spotted novel RBPs such as KhpA, a putative RNA chaperone, that potentially regulates sporulation.

## RESULTS

### Most putative independently transcribed sRNAs are specifically upregulated during sporulation

To assess the importance of RNA mediated regulation during sporulation of *B. subtilis*, we first used available transcriptomic data (Nicolas et al. 2012), from “tiling arrays” under numerous conditions that identified 1589 new RNA segments “of all categories” (antisense, UTRs, tRNAs, rRNAs, intergenic…). Among them, about 150 are independently expressed, making them putative sRNAs. We represented as a heatmap, the normalized expression profile of each of these independent transcripts during vegetative growth (in LB medium) and along sporulation (Fig. 2, Table S1). We observed than more than half independently transcribed RNAs are upregulated during sporulation, making this condition, among all those tested in (Nicolas et al. 2012), in which the largest number of sRNAs is overexpressed. We then looked for predicted dependencies on σ factors (Fig. 2 right column, Table S1) (Sierro et al. 2008; Nicolas et al. 2012). Consistently with experimental data, 66 transcripts (out of 152) exhibited dependencies on sporulation-specific σ^E^, σ^F^, σ^G^, σ^K^. Those putative sRNAs show none or very low expression during vegetative growth in LB. Notably, even if some non-coding RNAs do not appear specific from sporulation-specific σ, they are still enriched during sporulation process, suggesting their function is needed during the process, even if probably not specific to spore formation.

**Fig. 2:**
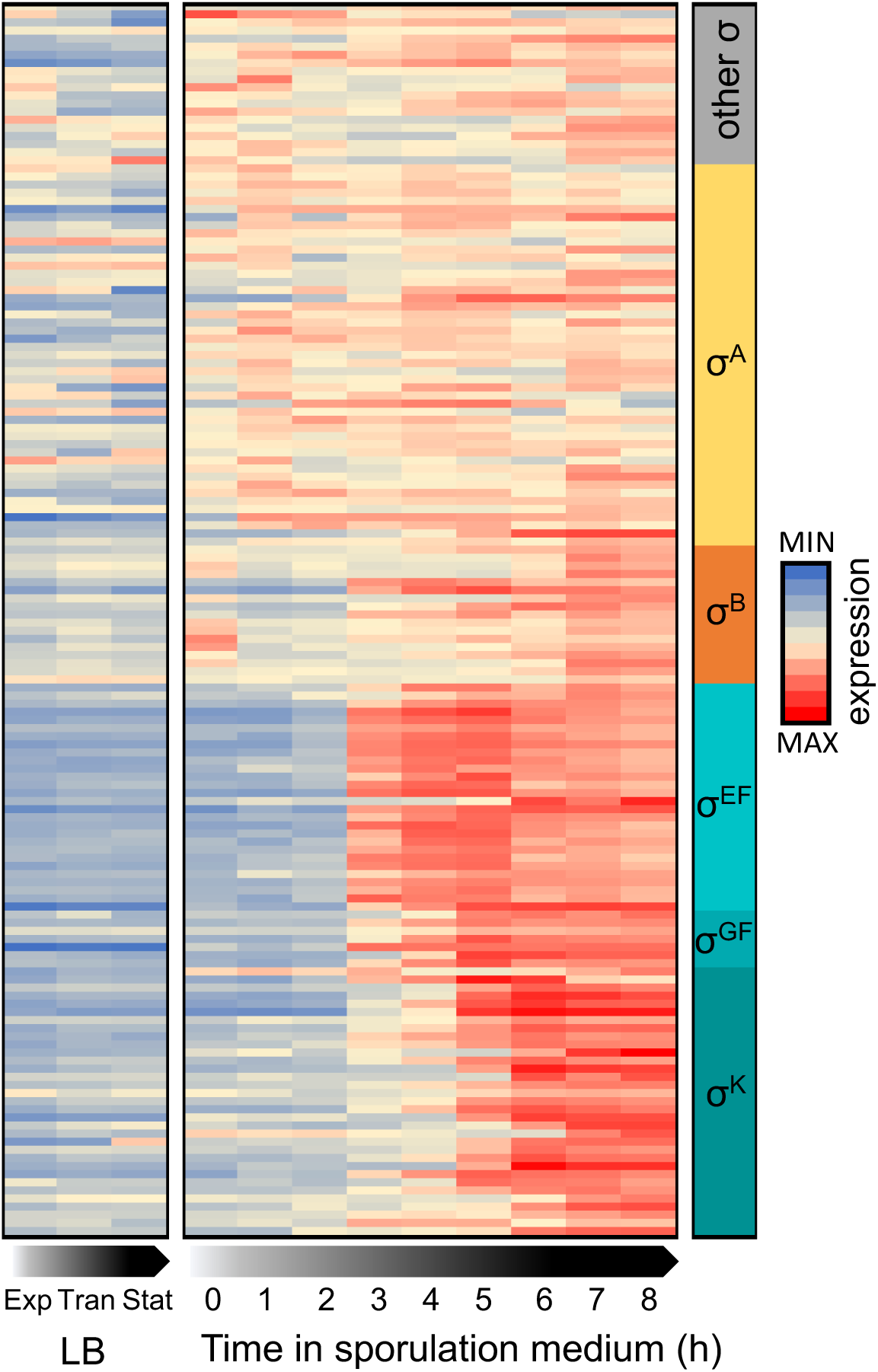
Heatmap showing normalized expression of sRNAs in LB (exponential, transition and stationary phases) and along sporulation (0h to 8h) sorted by predicted or known dependency on σ factor(s) (showed in the right column); (Sierro et al. 2008; Nicolas et al. 2012). Expression at a time point is shown as a fraction of the total expression across all shown conditions (sum of normalized quantity across each line = 1). As expected, sRNAs dependent on σ^E^, σ^F^, σ^G^, σ^K^exhibit strongest expression during sporulation.

We observe that most sporulation specific putative sRNAs are expressed under σ^E^ and σ^F^, corresponding to the early stage of sporulation, or under σ^K^, in late mother cell (Fig. 1), between 4h and 7h growth in sporulation medium (Fig. 2). These data suggest an important regulatory role of sRNA during sporulation but did not give information concerning their protein partners.

### RNA-protein complexes stabilized by UV crosslink can be purified by Orthogonal Organic Phase Separation (OOPS) in *B. subtilis*

To identify RNA-protein complexes involved in sporulation, we first conducted a kinetic of sporulation to define time points to look for RNA partners. *B. subtilis* was grown in sporulation medium (DSM) and observed at different times by contrast phase microscopy (Fig. 3A). Sporulation, though not synchronized among cells, starts when nutrients are depleted. After 3h in DSM, only vegetative cells are present. A fraction of them is probably initiating sporulation. First pre-spores are appearing after 6-7h in DSM. After 7h in DSM, the population consists of both vegetative and sporulating cells. After 12h, late pre-spores and first mature spores are observed, their number increasing with time. Comparing to conditions from (Nicolas et al. 2012), we observed that in our experiments, sporulation is slightly delayed (Fig. 2 and 3A). Early stages of sporulation (σ^E^, σ^F^) are triggered between 4 and 5h in Fig. 2 while our observation in DSM shows first pre-spores around 7h. All together, these observations suggest that most of RNA mediated regulations of sporulation take place between 7 and 12h in DSM. We thus chose to purify and analyze the RNA-Binding proteome (RBPome) after 3h, 7h, 12h of growth in DSM. For that, we carried out OOPS (Orthogonal Organic Phase Separation), that has the advantage of requiring less starting material than the other similar approaches. This approach combines *in vivo* UV crosslinking to stabilize RNA-protein interactions and acidic guanidine phenol chloroform extraction to enrich crosslinked protein-RNA complexes (Fig. 3B). Then, the different fractions were analyzed by mass spectrometry (Fig 3C). Since this approach has never been used for Gram-positive bacteria that possess a strong cell wall, principally composed of peptidoglycan, we adapted the protocol to *B. subtilis.* We optimized lysis conditions, as an insufficient lysis would populate the interphase with cell wall remnants, and adjusted UV irradiation to get robust RBP recovery (See Material and Methods). To assess crosslink and lysis efficiencies, we purified interphase from cells subjected or not to UV and treated it with protease instead of RNase. RNAs were recovered intact from the interphase (Fig. 4A) and quantified (Fig. 4B). We observed that the amount of RNAs present at the interphase of the non-crosslinked samples represents less than 5 % of total RNAs which is consistent with an efficient lysis (Villanueva et al. 2020). UV crosslinking increases protein-bound RNA recovery to reach around 20% of total RNAs (Fig. 4B). Notably, even without UV, RBPs are recovered at the interphase. Proteins isolated from the interphase (“RBPs”) and from the organic phase (“Free proteins”) were analysis by mass spectrometry followed by label-free quantification.

**Fig. 3:**
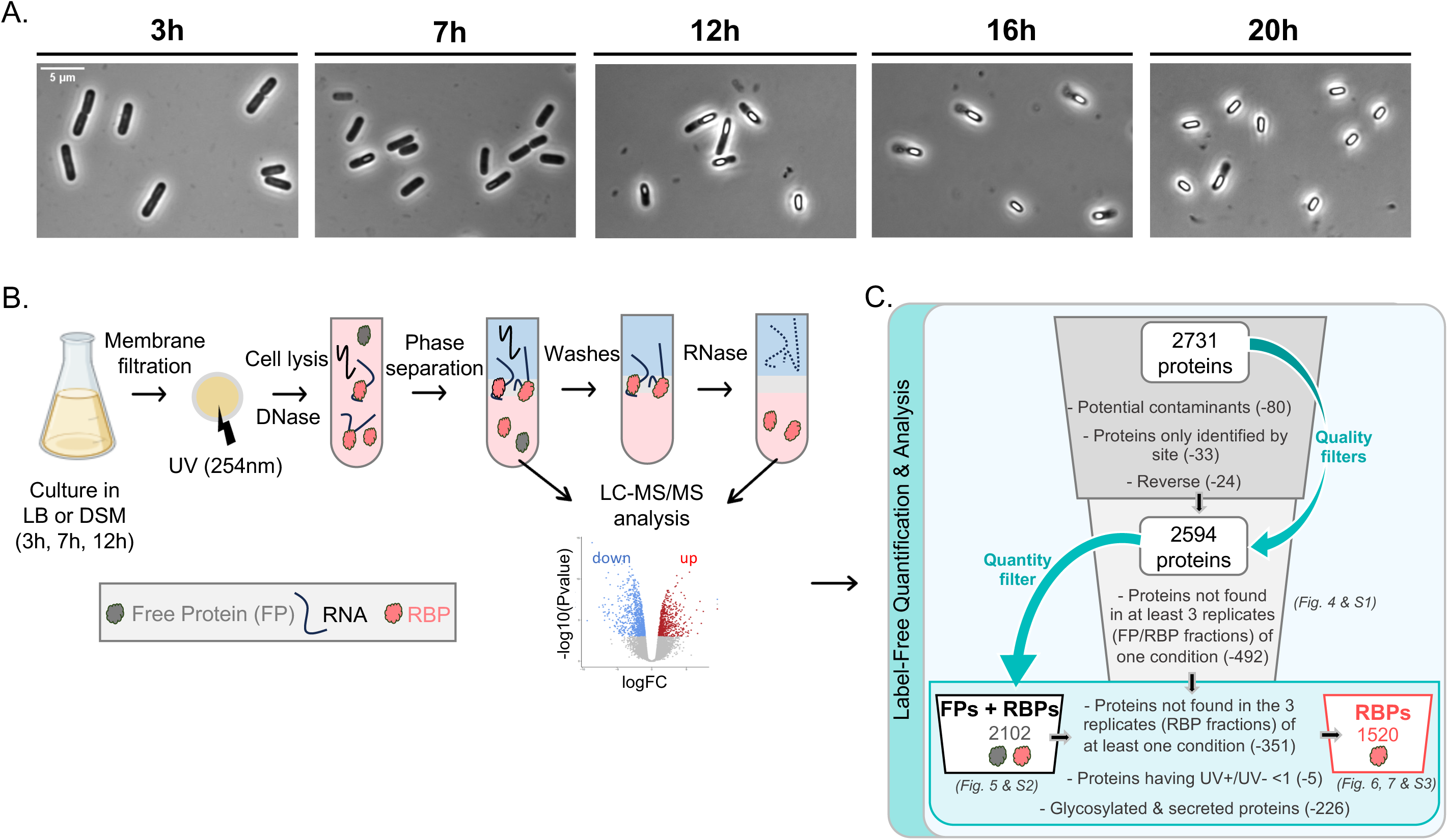
A. Observation in phase contrast microscopy of *B. subtilis* cells after 3h, 7h, 12h, 16h and 20h growth in DSM. B. Overview of OOPS protocol. To purify the whole RBPome involved in sporulation, we carried out OOPS at 3h, 7h, 12h in DSM. This approach combines *in vivo* UV crosslinking to stabilize RNA-protein interactions and acidic guanidine phenol chloroform extraction (AGPC, known as TRIzol) to enrich crosslinked protein-RNA complexes. After culture, cells are rapidly filtered and subjected to UVC crosslinking and lysed in presence of DNases. Lysates are then phase separated using TRIzol. Free RNAs go in the upper aqueous phase, free proteins in the lower organic phase and crosslinked RNA-protein complexes sharing properties of both RNAs and proteins in the interphase. After isolation, the interphase is enriched in crosslinked RNA-RBP complexes by several rounds of separation (washes) then treated with RNase and subjected twice to phase separation in order to recover only the RBPs. Free RBPs are then purified from the organic phase. Both free proteins (FPs) and RBPs are analyzed by liquid-chromatography mass spectrometry (LC-MS) and quantified by label free quantification (LFQ). C. Filters used for proteomic data analysis. Using MaxQuant Perseus software, 2731 proteins were listed in all the samples analysed. After utilisation of quality filters (potential contaminants, proteins only identified by site, reverse), 2594 proteins left. Then data were filtered to preserve only the proteins identified in at least three samples from one of the replicates (2102 proteins left corresponding to both free proteins and RBPs). Finally, RBPs were defined as the proteins once again identified in at least three replicates of a condition within the RBP fraction (1751 proteins left). Then we filtered out proteins showing a UV+/UV-ratio lower than 1 (1746 proteins left) and proteins annotated as secreted and glycoproteins that could be found in the interphase non-specifically (1520 RBPs left).

**Fig. 4:**
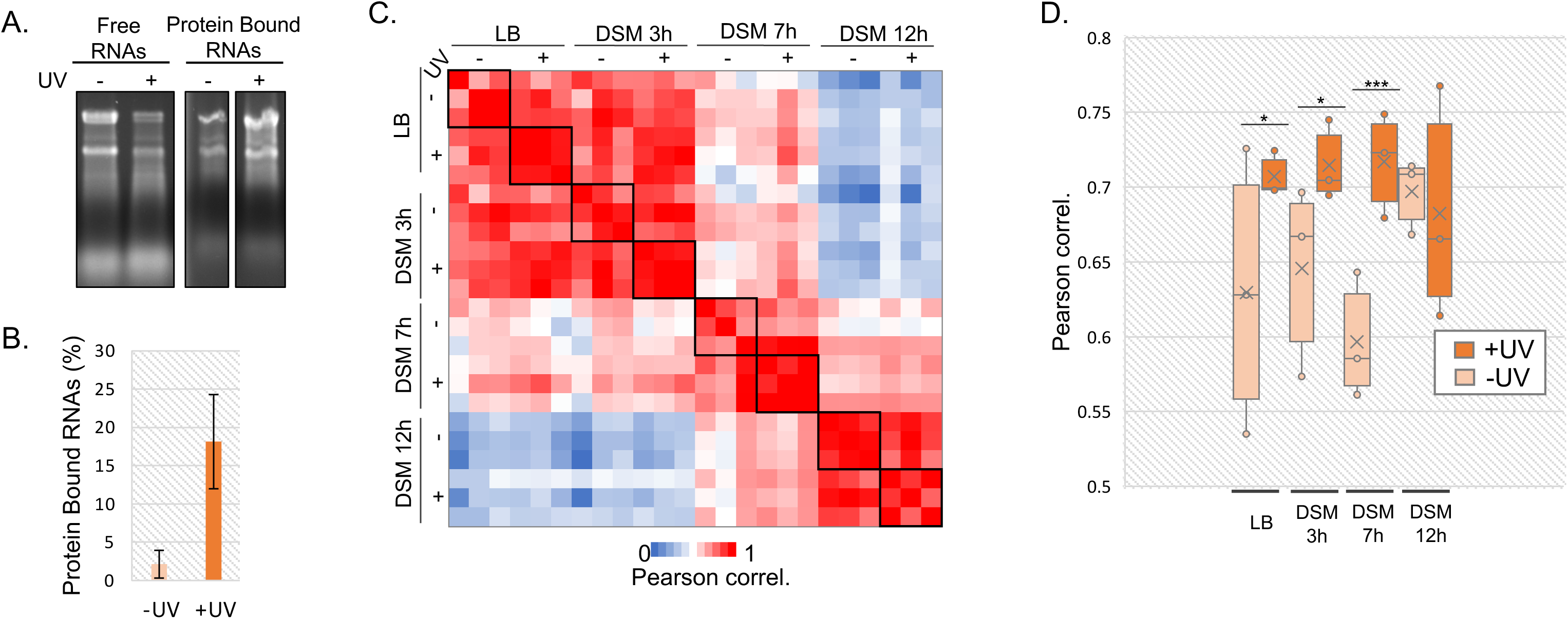
A. 1/20^th^ of quantity of free RNAs and 1/3^rd^ of quantity of proteins bound RNAs isolated from aqueous phase and interphase respectively, were separated on a 1.5% agarose gel and ethidium bromide stained. B. The proportion of isolated protein-bound RNAs with and without UV irradiation was calculated and displayed as a histogram. C. Correlation matrix displayed as a heat map showing Pearson correlation between datasets (with LFQ intensities) obtained for each individual replicates of RBP fractions in LB and along sporulation process in DSM medium. D. Distribution of Pearson correlation coefficient within triplicates for each growth condition. Higher coefficients indicate greater similarity between biological replicates.

To assess reproducibility and robustness of the approach, we calculated Pearson correlation coefficient to compare label-free quantification scores (LFQ) of proteins obtained in the three replicates of different fractions (Fig. 4C and S1A, Table S2) and carried out principal component analyzes (PCA) (Fig. S1BC). In the obtained correlation matrix, we observed highest correlation for RBPome (Fig. 4C) and free proteome (Fig. S1A) recovery between cells grown 3h in LB or DSM medium where cells are still in vegetative state (Fig. 3A). However, RBPome is more different after 7h or 12h growth in DSM, when sporulation has started (Fig. 3A).

PCA confirmed high reproducibility of RBPome recovery between replicates in each condition (Fig S1B). Replicates of free proteome exhibited more variability, especially for samples grown 7h in DSM (Fig S1C). As observed in the Pearson correlation matrix, PCA exhibits biological triplicates of LB and DSM 3h samples that are really close to each other while datasets obtained after 7h and 12h DSM samples are clustered independently (Fig. S1BC). Moreover, as expected, the RBP fraction exhibited low correlation with the free protein fraction (Fig. S1A), consistent with specific isolation of RBPs in the interphase.

Although UV does not appear to affect the free protein recovery (Fig. S1A, Table S2), UV crosslinked RBP fractions exhibit higher correlation than non-UV irradiated treated samples except after 12h growth in DSM (Fig. 4CD). To assess the effect of UV, we represented the distribution of correlation coefficients between replicates within a condition as a box-plot (Fig. 4D). Correlations between replicates are significantly higher for irradiated samples, except after 12h growth in DSM. Indeed, as we observed by microscopy (Fig 3A), mature spores are present at 12h and they are probably less or unaffected by crosslinking since one of the characteristics of spores is UV resistance, introducing some variability between datasets. Even with limited efficiency (Fig. 4B), UV crosslinking allows to recover higher quantity of RBPs with highest reproducibility between replicates.

### OOPS allows the specific purification of RBPs along sporulation

To confirm the specificity of the method to isolate proteins with RNA binding ability, we checked if the recovery of a protein in the RBP fraction depended solely on its RNA binding ability and not on its production level. We thus compared the distribution of relative abundance in RBP and “free protein” fractions. Proteins were sorted by relative abundance in free protein fraction and their LFQ intensities were displayed as a heat map (Fig. 5A and S2A, Table S3). We observed that distributions of LFQ intensities are not correlated between both fractions. Some highly produced proteins exhibit low LFQ in RBP fraction while many low produced proteins are highly recovered in RBP fraction. The recovery of a protein in the RBP fraction is thus independent of its production level.

**Fig. 5:**
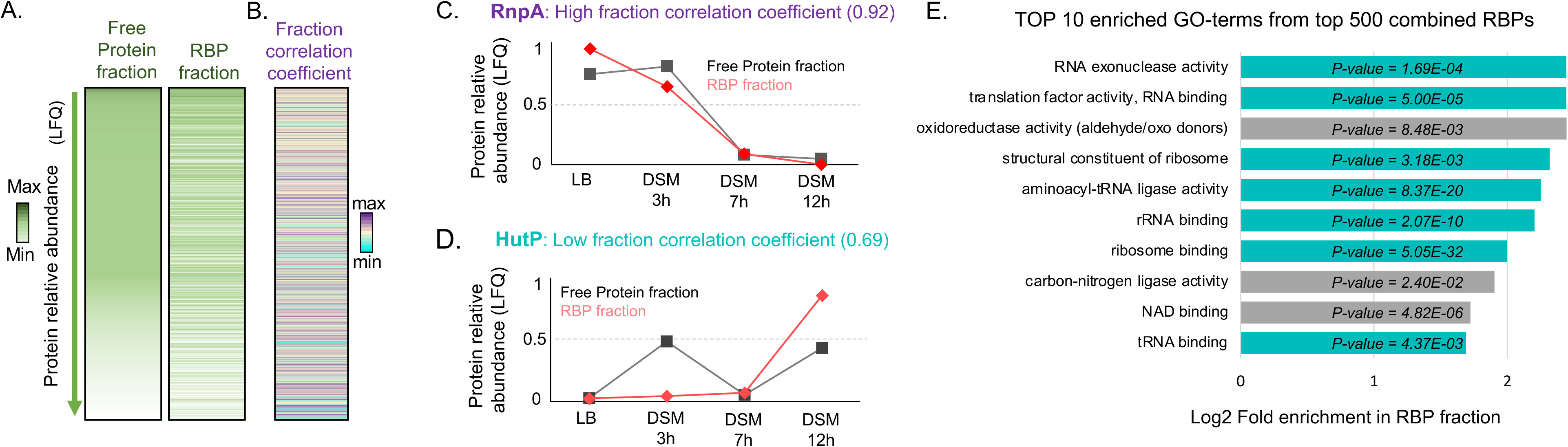
A. Heatmap showing protein abundance (LFQ) in both free protein and RBP fractions (within each fraction, LFQ intensities from all 4 conditions are summed up for each protein). B. Corresponding fraction correlation coefficient. Heatmap is sorted by average abundance in free protein fraction. A high fraction correlation coefficient corresponds to proteins always bound to RNA, their abundance profiles in both fractions are similar. A low fraction correlation coefficient corresponds to proteins transitory bound to RNA; they exhibit different profiles in both fractions. All values are displayed in Table S3. Representation of normalized abundance profiles in both free protein and RBP fractions for RnpA, a protein always bound to RNA, exhibiting a high dynamic correlation coefficient (C.) and HutP, a protein bound to RNA only in certain conditions (D.). E. Top 10 enriched GO-terms from the top 500 RBPs from the different conditions (all p-values < 0.025).

Furthermore, to assess if RNA binding profile along sporulation was dependent on its production, we calculated, for each protein, a fraction correlation coefficient (Fig. 5B, Table S3), *i.e.,* the correlation coefficient between profiles of recovery in both fractions (See Material and Methods). This indicates how different is the relative recovery at each point between the two fractions. All fraction coefficients in our data are between 0.29 and 1 (median = 0.82) (Fig. S2B). For RBPs always bound to RNA, we expect their recovery in RBP fraction to be following their production level, thus exhibiting a high correlation (close to 1) between their modulations in RBP fraction and in free proteome. This is illustrated by two proteins that constitutively bind RNA: RpoE (Fig. 5C), a subunit of the RNA polymerase, and RnpA, a protein component of RNase P that have fraction correlation coefficients of 0.94 and 0.92, respectively (Table S3). On the contrary, we expect that proteins binding RNA transitory will have lower fraction correlation coefficient and RNA binding abilities (recovery in RBP fraction) modulated independently from their production level (recovery in free protein fraction). For example, the anti-terminator HutP, regulating the *hut* operon by binding a terminator RNA structure in the presence of histidine (Oda et al. 2004) (Fig. 4D), exhibits a low fraction correlation coefficient (0.69). It is the case of other anti-terminators such as LicT (0.77) (Declerck et al. 1999), SacT (0.71) (Arnaud et al. 1996) and RplT (0.78) (Deiorio-Haggar et al. 2013). All these proteins bind RNA only in specific conditions and have coefficients lower than 0.82 corresponding to the median value (Table S3). Notably, we observed that the fraction correlation coefficient is, as expected, independent of RBP abundance (Fig. 5AB and S2C).

The top 500 proteins identified in the RBP fractions (from all samples combined) was analyzed for Gene-Ontology (GO) terms enrichment (Fig. 5E). Among the 10 most enriched GO-terms, 7 are related to RNA function. Remarkably, the 3 remaining enriched GO-terms are related to metabolic enzymes, consistent with the observations made in eukaryotic organisms and in *E. coli* and *S. aureus* (Chu et al. 2022; Urdaneta et al. 2019; Trendel et al. 2019; Shchepachev et al. 2019) suggesting these enzymes would moonlight as RBPs to link directly the response to nutrient availably in the environment with regulation of gene expression. Notably, the well-known moonlighting protein CitB, both metabolic enzyme and RBP in iron starvation conditions (Pechter et al. 2013), whose deletion results in a strong sporulation defect (Craig et al. 1997), was found in the top 100 RBPs isolated after growth in LB and DSM (3h, 7h, 12h).

### RBPome is highly remodeled during sporulation process

Among the 2102 proteins from both free protein and RBP fractions, 1520 passed the filters to be considered as putative RBPs (Fig. 3C): being identified in at least 3 biological replicates of one condition of the RBP fractions, exhibiting enrichment in crosslinked samples compared to the non-crosslinked samples and not known as being secreted and/or binding or interacting with glycans (including spore crust proteins). Indeed, glycoproteins could be not specifically found in the interphase (Queiroz et al. 2019; Esteban-Serna et al. 2023).

We then looked how specific these datasets were for a condition (Fig. 6, Table S4). 70% of proteins (1049/1520) were recovered in RBP fractions across all 4 conditions; among them, almost all (54/60) ribosomal proteins encoded in the genome. Interestingly only 2% of proteins seem to be exclusive to the vegetative state (LB and or DSM 3h) while more proteins were recovered specifically during sporulation process. Indeed, more than 13 % RBPs appear specific of DSM 7h and 12h (either exclusively or in both time points). This indicates that RBPs exhibit different RNA binding and production profiles across sporulation.

**Fig. 6:**
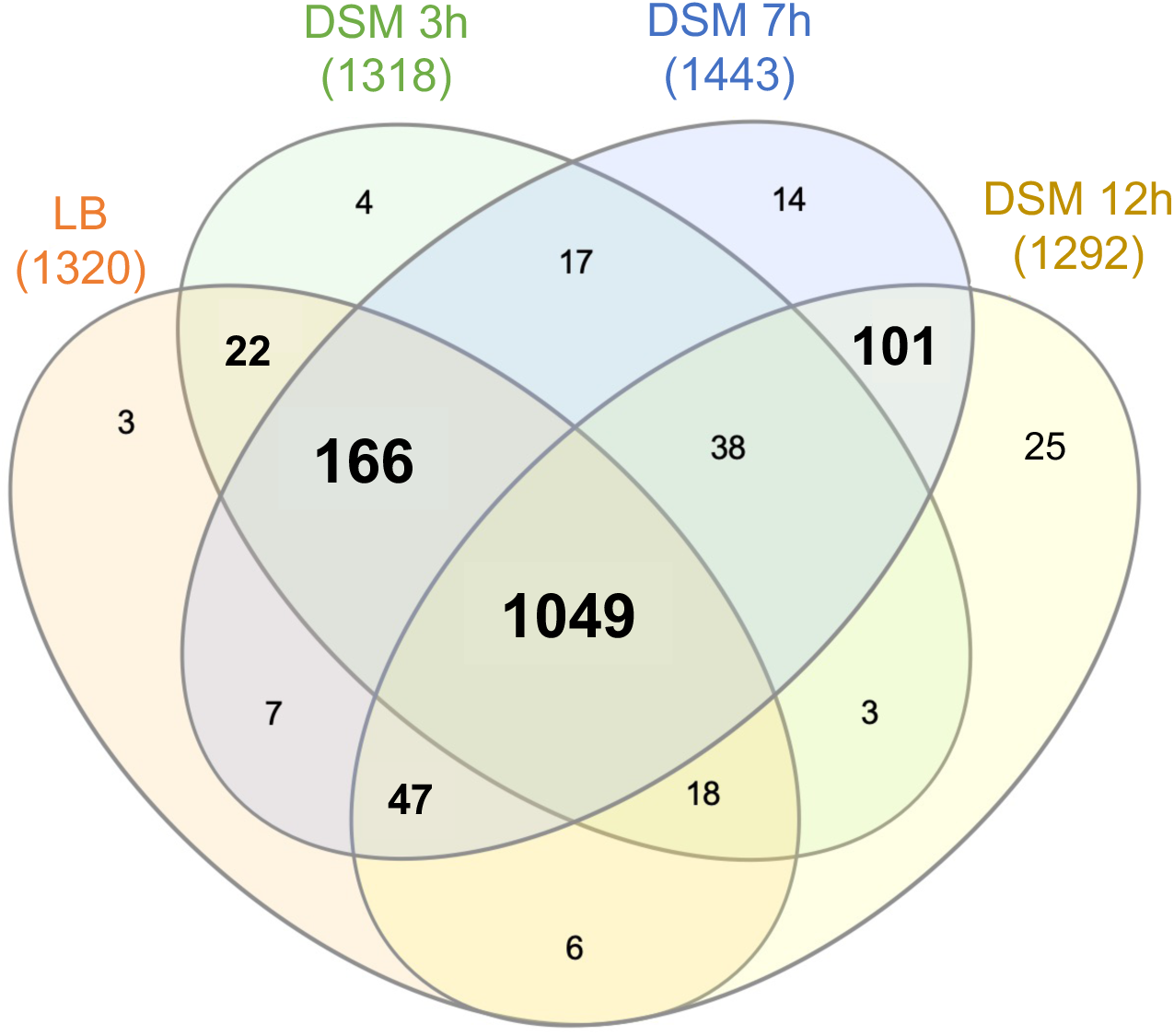
Venn diagram showing the overlaps between RBPs identified in cells grown in LB and after 3h, 7h and 12h in DSM.

### Classification of RBPome in different clusters

To compare both production and RNA binding ability of RBPs, we carried out clustering analysis (based on Euclidian distance) to group together proteins exhibiting similar behavior (Fig. 7, Table S5). In each cluster, proteins known to be an RBP, known or predicted to have a role in sporulation, in metabolism or to be essential were highlighted. In addition, GO-term enrichments were explored for each cluster of RBPs. Clustering analysis has been performed on data in which intensities were normalized per gene (where recovery score at each condition is the fraction of the total recovery of this protein in a given fraction) and distribution of unnormalized LFQ values in each fraction was considered (Fig. S3).

- In **Cluster 1,** proteins exhibit similar profile in both fractions, with exclusive high recovery in LB. Most of them are still present after 3h growth in DSM but don’t appear to bind RNA anymore. No GO-term were enriched.
- **Clusters 2 and 3** seem composed of the most abundant proteins in the cells, specific from vegetative growth, exhibiting highest production level after 3h growth in DSM but maximum RNA binding in LB. Their synthesis shut down after 7h and 12h in DSM. We observed significative enrichment for proteins involved in respiration (menaquinone biosynthesis), transcription (RNA biosynthesis), metabolism (pentose-phosphate pathway). Proteins involved in division and chemotaxis are enriched too. Altogether, these observations are consistent with the cells being gradually less metabolically active through sporulation process. Indeed, both translation and transcription machineries have been shown to be slowed down during sporulation process (Iwańska et al. 2023), consistent with the profiles obtained at 7h and 12h in DSM. SpoVG, protein with a potential role in sRNA processing (Christopoulou and Granneman 2022) was found in cluster 3. 12 RNases and RNAse cofactors were recovered in clusters 2 and 3 (Table S6).
- **Cluster 4** is composed of abundant proteins (Fig. S3), enriched in metabolic enzymes and proteins involved in resistance with oxidative stress and toxic substances. Interestingly most of them are specifically highly produced after 3h in DSM. Nevertheless, they are almost only found in the RBP fraction after 12h in DSM suggesting they bind RNAs under nutrient starvations or stresses. Indeed, in stationary phase, toxic metabolites accumulate in growth medium; genes involved in protecting the cells against stresses such as oxidants and toxic chemicals are expressed (Nyström 2004; Dukan and Nyström 1999). 32 / 231 proteins found in cluster 4 are known to bind RNA, among which Hfq. Notably, Jag, which was not identified yet in *B. subtilis* as an RBP, has sRNA binding ability in other bacterial species and was recovered in this cluster (Zhu et al. 2024).
- As cluster 4, proteins of **cluster 5** exhibit low fraction correlation profiles, their production reaching highest level after 3h and 12h in DSM but found in RBP fraction only after 12h growth in DSM. They must transitorily bind RNA during sporulation. No GO-term was enriched for this cluster although lots of metabolism proteins, that could be moonlighting as RBPs, are present. It is worth noting that proteins are present in low quantity in both fractions, which likely contributes to noisy recovery.
- **Cluster 6** is enriched in rRNA metabolism enzymes consistent with a permanent RNA binding activity. Most proteins from cluster 6 are not highly produced but they are recovered in higher quantity in the RBP fractions suggesting they strongly bind RNA (Fig. S3). Most of them exhibit high correlation between production profile and RNA binding function. Genes involved in amino-acids metabolism are enriched (36/177). Besides being proteins building blocks, amino-acids serve for metabolism and survival (Dai et al. 2022).
- RBPs from **cluster 7** are only observed after 3 h in DSM and their RNA binding ability is highly correlating with their production profile, suggesting a fraction of the produced proteins binds RNA. They are indeed less recovered in the RBP fraction than the free protein fraction (Fig. S3).
- RBPs found in **clusters 8, 9 and 10** are highly enriched in sporulation proteins and exhibit the same profiles between both fractions (cluster 8: 29/58 RBPs; cluster 9: 48/122; cluster 10: 51/106). Cluster 8 is composed of RBPs mainly produced after 7 h in DSM while RBPs of cluster 9 are found both at 7 h and 12 h growth in DSM. Their abundances remain relatively low compared to RBPs from other clusters, but are higher in the RBP fraction than the free protein fraction. This suggests that most proteins of this cluster are bound to RNA. Notably factors from fatty acid metabolism pathway are highly enriched, consistent with the increase of lipid production required to the spore formation (Schujman et al. 1998; López et al. 1998). We propose that this pathway may use RNA mediated regulation. Within cluster 10, RBPs are specifically produced after 12h in DSM. This cluster is depleted for proteins involved in cellular processes, gene expression, translation and RNA metabolism, most cells being less metabolically active. Interestingly, proteins involved in isoleucine and serine, glutamine and arginine metabolism are recovered and enriched across several clusters (6, 8 and 9 respectively) suggesting an RNA mediated regulation of amino-acid metabolism.

**Fig. 7:**
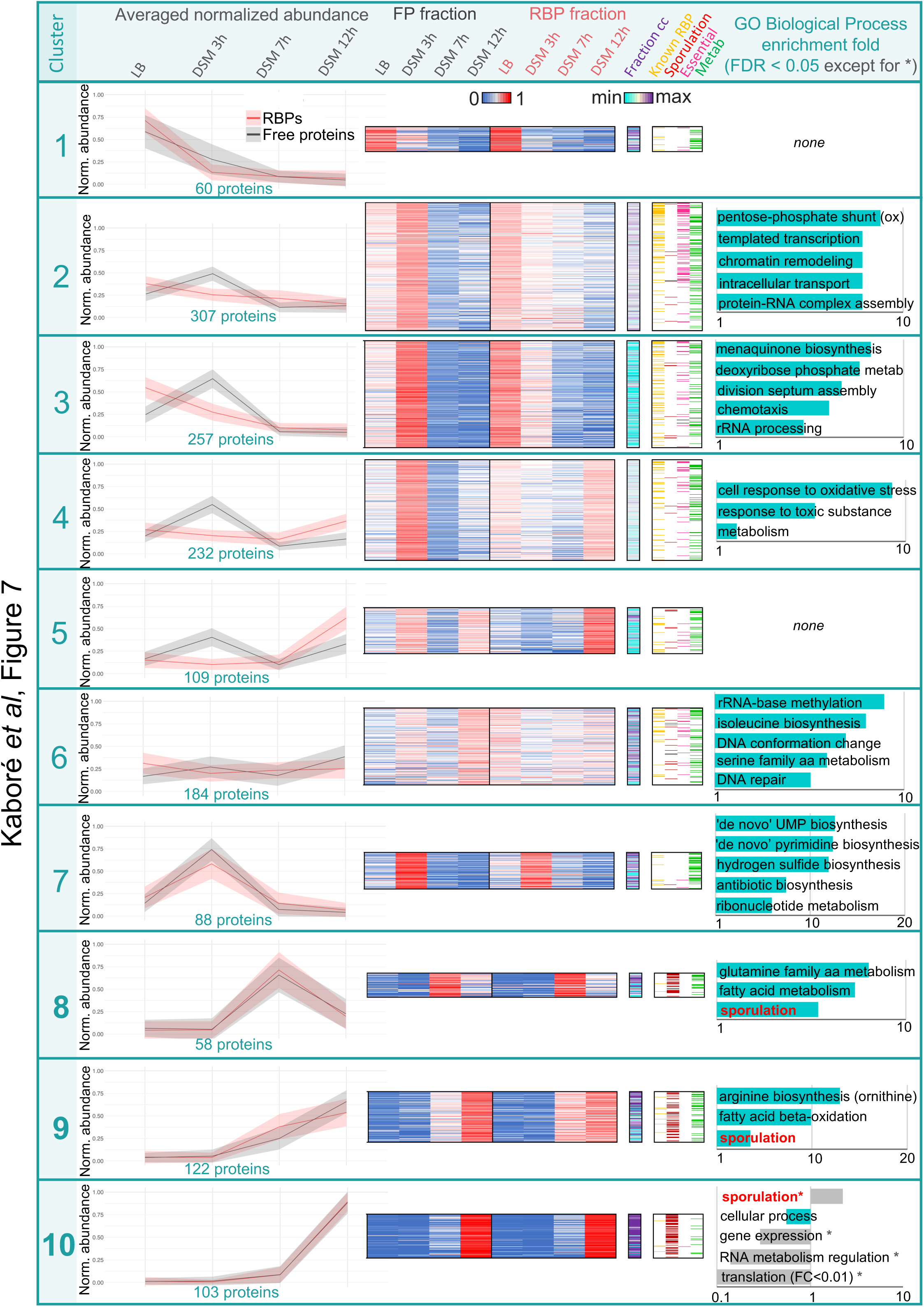
Normalized LFQ intensities in both free protein and RBP fractions (corresponding to production and RNA binding profiles respectively) displayed as a heatmap. Each line corresponds to a protein. For each condition, abundance is expressed as a fraction of the total recovery of this protein across all four tested conditions, either in the free protein (FP) or RBP fraction (sum of normalized quantity across each line = 1). Proteins have been clustered by Euclidian distance in 10 distinct groups according to their profiles. All values are displayed in Table S5.

### Highlight on unknown proteins binding RNA specifically during sporulation

Among identified RBPs by OOPS, we expect sRNA partners involved in setting up sporulation to harbor an RNA binding ability that fluctuates along sporulation. We consequently focused on RBPs whose presence significantly increased at 7h in DSM, conditions where more sRNAs are expressed and most cells are ongoing sporulation process, but are still not mature spores. We decided to focus on clusters 8 and 9 (Table S7) consisting of 180 proteins (including 83 proteins known as involved in sporulation).

Among the 180 proteins, 11 are potential or characterized RNA binders (*spoIID, kipR, pksL, helD, sigF, sigK, sigE, sigG, mfd, rnr, nrnB*). As noted in the global results, a significant part of clusters 7 and 8 carries enzymatic activities linked to cell metabolism, being probably moonlighting enzymes -those exhibiting two or more physiologically relevant biochemical or biophysical functions (Chen et al. 2021) (Curtis and Jeffery 2021). 45 proteins in the selection have uncharacterized function even if most of them (38/45) are referenced as putative enzymes.

We decided to concentrate our interest on the 7 other unknown proteins, listed in Table 1, to assess whether they could be partners of non-coding RNAs, looking for conservation, structural homology, their σ factor dependency, and data from literature and the *Subtiwiki* database (Zhu and Stülke 2018). These different proteins exhibit diverse recovery scores (free protein fraction) and enrichment in the RBP fraction. They are all conserved in other spore-forming firmicutes, except YxnB, which would be consistent with a role in sporulation. Interestingly, among these 7 proteins, is KhpA (YlqC), a highly conserved putative RNA chaperone uncharacterized in *B. subtilis*.

**Table 1:**
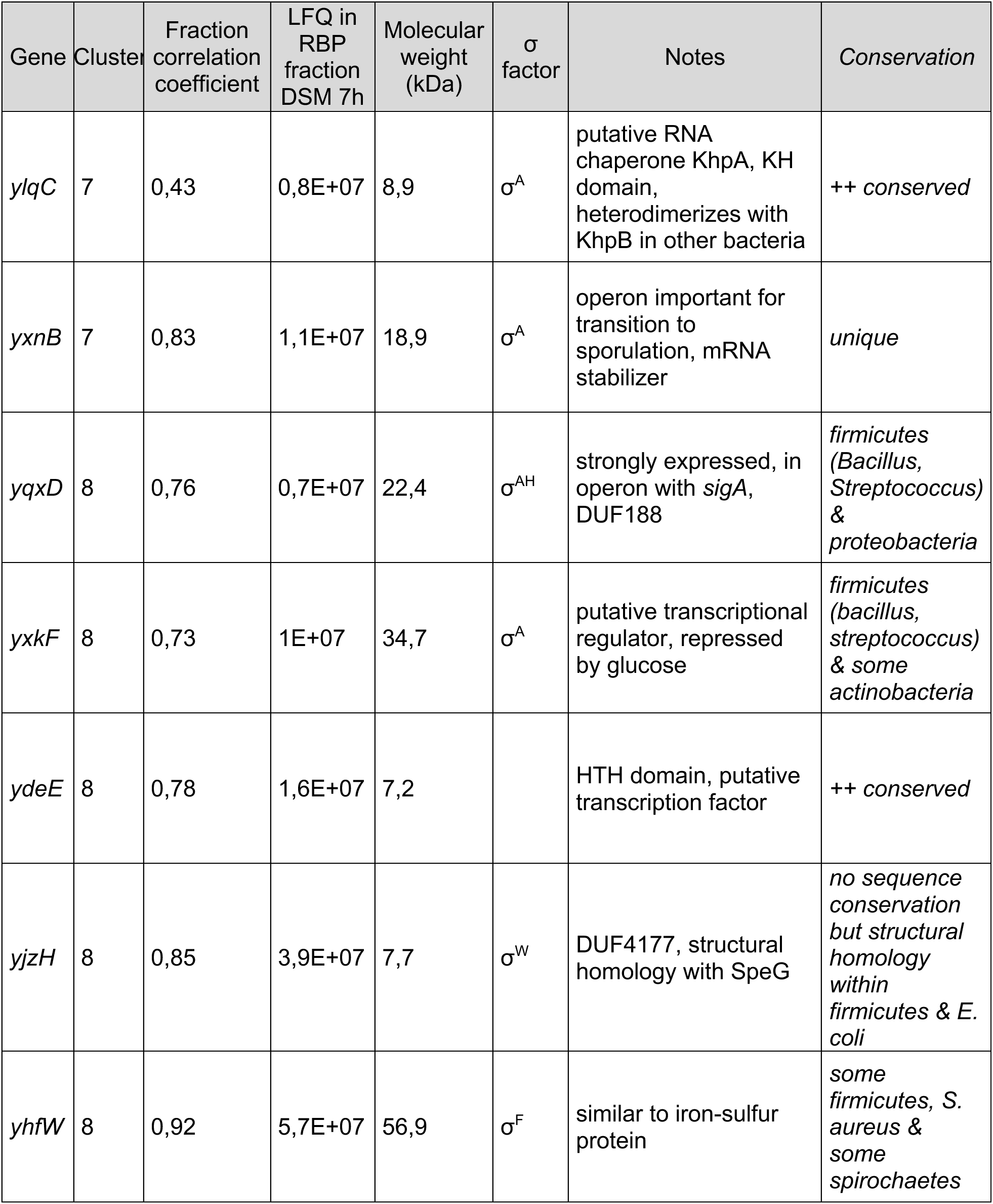
Proteins of unknown function recovered in clusters 7 and 8.

The *yxnB* gene belongs to an operon involved in adaptation to nutrient starvation and transition to sporulation. It is unique in databases and its function is unknown but its presence in the operon transcript has been shown to stabilize the downstream part of the operon mRNA (Morinaga et al. 2010).

*yqxD* is part of the operon that contains genes encoding a DNA primase involved in the chromosome replication and σ^A^. It was shown to be dispensable for vegetative growth but affecting sporulation initiation (Zuberi and Doi 1990). YqxD has a PIN-like domain known to be involved in ribonuclease activity (Matelska et al. 2017).

YxkF and YdeE are putative transcriptional regulators with Helix-Turn-Helix (HTH) domains, overexpressed during sporulation.

YjzH is a small DUF4177 domain-containing protein overproduced under glucose exhaustion (Nicolas et al. 2012). Its sequence is poorly conserved but we found it has a strong structural similarity (Hutson 2023) with SpeG, a conserved polyamine acetyltransferase involved in biofilm formation in other *Bacilli* which did not have homolog identified in *B. subtilis* yet (Tsimbalyuk et al. 2020). It was shown to regulate expression of several genes (Fang et al. 2017), such as the sRNA rprA in *E. coli* (Hu et al. 2018).

YhfW is an iron-sulfur putative oxidoreductase, strongly upregulated in sporulation and competence affecting transformation efficiency, sporulation and germination through uncharacterized mechanisms (Boonstra et al. 2020).

Altogether, our results show that these proteins bind RNAs during sporulation process leading to new hypothesis suggesting they would be involved in RNA-mediated regulation.

### KhpA, an RBP with potential role in sporulation regulation

We further investigated the role of KhpA in sporulation. Among the sporulation-specific clusters (8, 9, and 10), KhpA exhibited the lowest fraction correlation coefficient (0.43), suggesting that it does not always bind RNA. KhpA is highly abundant in vegetative cells and its production decreases during sporulation (Fig. 8A) but, it is detected in the RBP fraction only after 7 hours in DSM. This suggests that KhpA specifically binds RNA during early sporulation, despite being less expressed than in vegetative cells. To explore KhpA RNA-binding activity, a strain expressing KhpA fused to a 6xHistidines and FLAG domain (HF tag) under the control of its native promoter was constructed. After growth in LB to transition phase and DSM for 7h, cells were subjected to UV crosslinking. KhpA-HF was then tandem-affinity purified under both native and denaturing conditions, ensuring high stringency in the recovery of authentic RNA-protein interaction sites. Following ^32^Phosphate labelling and gel separation, KhpA-associated RNAs were detected (Fig. 8B), both under sporulation and, to a lesser extent, vegetative growth, although no signal was detected in OOPS.

**Fig. 8:**
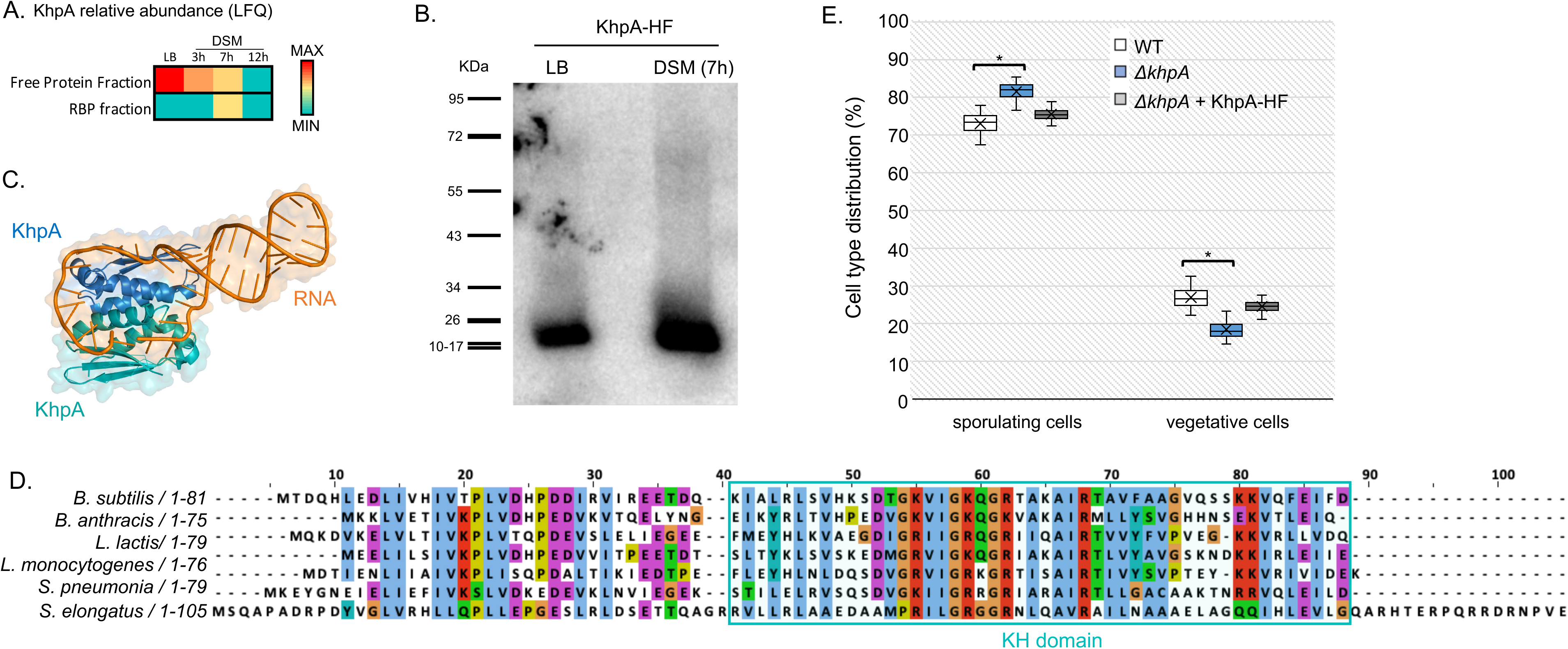
A. KhpA relative abundance (LFQ) in OOPS free protein and RBP fractions, along sporulation. B. *In vivo* RNA binding assay on KhpA-HF in LB (transition phase) and DSM (7h). ^32^P labeled RNAs crosslinked to KhpA-HF were visualized using a PhosphorImager. C. Structure of KhpA homodimer in complex with a poly-U RNA, predicted by Alphafold 3 (Abramson et al. 2024). D. Sequence and domain alignments of KhpA from different bacteria (*B. subtilis, B. anthracis, Lactococcus lactis, Listeria monocytogenes, Streptococcus pneumonia, Synochococcus elongatus)*. KH domain is highlighted in blue; its minimal structure is characterized by the presence of two α-helices flanking a β-sheet on each side, as well as a GxxG motif between the two α-helices. This motif, along with the amino acids of the adjacent β-sheet, forms an RNA-binding surface. E. Sporulation efficiency assay; fraction of sporulating cells (spores + prespores) is calculated relative to total cells (vegetative + prespores + spores) after 24h in DSM. Tukey tests were performed and showed significative differences between WT and *ΔkhpA* strains (* = p-values < 0.05). Differences between *ΔkhpA* and complemented strains exhibit p-values < 0.01 for both cell types.

The majority of the KhpA sequence consists of a KH domain, a well-characterized RNA-binding motif, highly conserved among bacteria (Fig. 8CD). KhpA has been previously reported to interact with RNAs, including sRNA, in *Streptococcus pneumoniae* or *Fusobacterium nucleatum* (Lamm-Schmidt et al. 2021; Michaux et al. 2023; Zhu et al. 2024) and was shown to be able to form homodimer in *S. pneumoniae* (Winther et al. 2019). Using Alphafold3 (Abramson et al. 2024), structure of a KhpA homodimer in contact with an RNA (polyU in that case) was predicted. Without RNA, the confidence score (interface predicted template modelling or ipTM) is really low (0.35) suggesting likely a failed prediction but the presence of the RNA increases the ipTM to 0.65 compared to the dimer alone. This suggests that the RNA would stabilize the protein-protein interaction that would be unlikely in its absence. To assess KhpA’s potential role in sporulation, we analysed a strain lacking *khpA*. The deletion strain exhibited a slight but reproducible increase in sporulation efficiency, complemented by expression of *khpA* under the control of its native promoter, suggesting a weak regulatory role in this process (Fig. 8E).

### SpoVR, a protein affecting sporulation with potential RNA-mediated regulation

Within cluster 8, SpoVR, a protein involved in spore cortex formation and whose absence induces a decrease in spores resistance to heat (Beall and Moran 1994), caught our attention. SpoVR is a protein conserved in many spore-forming bacteria from *bacilli* to gram-negative *myxococcales*. It was shown to be involved in spore cortex formation in *B. subtilis* (Beall and Moran 1994) and *Myxococcus xanthus* (Tengra et al. 2006). It was recovered specifically after 7h and 12h in DSM in both free protein and RBP fractions with highest RNA binding ability after 7h in DSM (Fig. 9A). To confirm RNA binding ability, an *in vivo* RNA binding assay was performed (as precedingly for KhpA) on a strain expressing SpoVR-HF under the control of its σ^E^ dependent promoter, during both vegetative growth (LB, transition phase) and sporulation (7h in DSM). ^32^P labeled RNAs crosslinked to SpoVR-HF were detected as expected during sporulation. Interestingly, an Alphafold3-predicted structure of SpoVR – poly-U RNA was obtained despite the absence of a known RNA recognition domain.

**Fig. 9:**
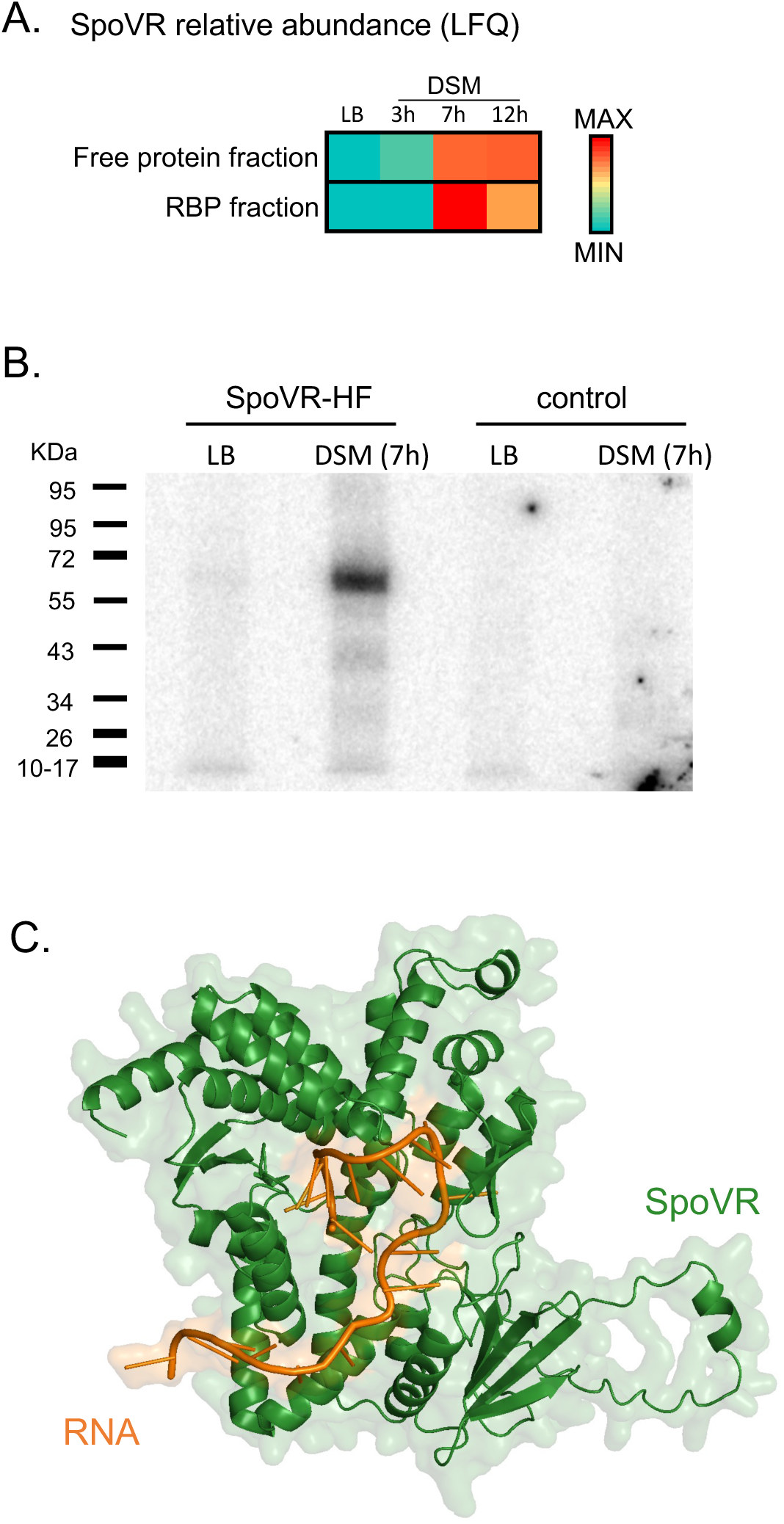
A. SpoVR relative abundance (LFQ) in OOPS free protein and RBP fractions, along sporulation. B. *In vivo* RNA binding assay on SpoVR-HF in LB (transition phase) and DSM (7h). ^32^P labeled RNAs crosslinked to SpoVR-HF were visualized using a PhosphorImager. C. Structure of SpoVR in complex with a 20-U RNA, predicted by Alphafold 3 (Abramson et al. 2024).

## DISCUSSION

In this paper, we followed the evolution of RNA-Binding Proteome during sporulation by OOPS, a powerful technique for specific purification of RBPs. By applying the OOPS technique for the first time to a Gram-positive bacterium, we have gained a comprehensive overview of RBPome during both vegetative growth and sporulation of *B. subtilis*. As proof of concept, we found that more than 230 known RNA-associated factors were robustly identified in the RBP fractions, with additional proteins characterized or suggested in other bacteria as sRNA-interacting factors. These include YvcJ, a homolog of *E. coli* RapZ (Cluster 2), an sRNA-binding RNase adaptor (Durica-Mitic et al. 2020; Khan et al. 2020), the RNA chaperones Hfq (cluster 4) and SpoVG (cluster 3) (Christopoulou and Granneman 2022), Kre (YkyB) in cluster 1 (Gamba et al. 2015), KhpA in cluster 8 (Lamm-Schmidt et al. 2021; Michaux et al. 2023; Zhu et al. 2024) and Jag in cluster 4 (Lamm-Schmidt et al. 2021; Zhu et al. 2024; Michaux et al. 2023). Additionally, RNA chaperones such as CspB, CspC, and CspD - preventing the formation of mRNA secondary structures to facilitate translation under stress conditions (Zhang et al. 2018; Faßhauer et al. 2021) - were identified in clusters 2 and 3 in the RBP fraction from vegetative cells. The recovery of these RBPs confirms that our experimental conditions allow the identification of proteins that act as post-transcriptional regulators through specific sRNA binding. 66 proteins containing an Sm domain, involved in protein oligomerization and RNA binding, (out of the 116 encoded in B*. subtilis* genome) were recovered in OOPS (Table S10). Sm (and Lsm (like-Sm) such as Hfq) containing proteins belong to a large family of proteins from all cellular organisms that are involved in numerous processes associated with RNA processing and gene expression regulation (Achsel et al. 2001).

Among the 30 known RNases and RNase cofactors in *B. subtilis*, 25 were identified using OOPS, exhibiting diverse recovery profiles in both the free protein and RBP fractions. This suggests process- and target-specific functionalities (Table S6). Notably, we identified KapD and nano-RNase B (NrnB), two exoribonucleases specifically expressed during sporulation. KapD was found in cluster 6, likely due to its basal RNA-binding activity in vegetative cells, whereas NrnB was identified in cluster 9, both displaying peak production and RNA-binding activity at late sporulation stages. This is consistent with previous studies showing that KapD is exclusively produced in the mother cell, where it regulates *sigK* mRNA stability, while NrnB functions in the forespore, degrading dinucleotides representing the final step of mRNA breakdown (D’Halluin et al. 2025; Myers et al. 2023). Additionally, RNases PH and R, although not specific to sporulation, were highly abundant during this process. Beyond RNases, three of the four RNA helicases encoded in the *B. subtilis* genome—CshA, CshB, and CshC (YfmL)— were identified among RBPs in clusters 3, 2, and 6, respectively. While CshA is linked to the RNA degradation machinery (Haq et al. 2024), the role of CshC remains unclear, although studies suggest its involvement in ribosome assembly and sporulation in *B. subtilis* (González-Gutiérrez et al. 2018). These proteins are conserved among *Bacilli* and are required for *B. cereus* sporulation, with CshC additionally contributing to oxidative stress resistance (Pandiani et al. 2011), likely playing key unelucidated roles in survival strategies.

Interestingly, numerous sporulation-specific factors were found to bind RNA, particularly at late sporulation stage (DSM 12h) like GerW and YutG, identified in cluster 10. Additionally, the presence of stable RNAs in mature spores (Setlow and Christie 2020; Zhou et al. 2023) and the large number of sRNAs controlled by the late sporulation-specific σ^G^ and σ^K^ (Fig. 2) suggest the existence of RBPs that stabilize these RNAs. However, technical challenges remain in studying these complexes, particularly in lysing spores while preserving RNA-RBP integrity and overcoming the resistance of mature spores to UV irradiation used for stabilization. Addressing these limitations will be essential for further investigations.

Technological advances in studying RNA-protein interactions have revealed a surprisingly high number of unconventional RBPs such as conserved metabolic enzymes (Caudron-Herger et al. 2021). Many of these proteins were primarily unrelated to RNA metabolism: among them, all enzymes of glycolysis and Krebs cycle (Beckmann et al. 2015; Hentze et al. 2018). While overlooked, multifunctionality in proteins is common. More than 500 proteins have been classified as moonlighting proteins. Beyond them, the aconitase CitB (Pechter et al. 2013) interacting with RNA, was recovered in the top 300 most abundant RBPs. Besides enzymes, 57 proteins containing HTH domain (out of the 245 encoded in the genome) were identified across the different conditions (Table S10). Usually considered as transcriptional regulators binding DNA, our results suggest that some of them can also bind RNA under specific conditions. This was already shown with CcpA, the master regulator of catabolite repression in several Gram-positive bacteria, able to bind hundreds of RNAs, to probably regulate their stability (Chu et al. 2022; Birkholz et al. 2024). Structural analyses have recently showed how an HTH domain can specifically discriminate between DNA and RNA (Birkholz et al. 2024).

Concerning KhpA, it has been previously reported to interact with RNAs, including sRNA, in complex with the protein KhpB in *S. pneumoniae* or *F. nucleatum* (Lamm-Schmidt et al. 2021; Michaux et al. 2023; Zhu et al. 2024). KhpB (also named Jag) was found among OOPS-isolated RBPs in cluster 4. Both KhpA and Jag contain RNA binding domains and have conserved homologs in Gram-positive bacteria (Olejniczak et al. 2022). Most research about KhpA and KhpB has been done in *S. pneumoniae*, where KhpA forms homodimers or heterodimers with KhpB. Strains lacking either one exactly phenocopy each other (Winther et al. 2019; Zheng et al. 2017) and have joint roles in growth and cell elongation. Our data suggest that KhpA could have specific role in sporulation in *B. subtilis* in absence of KhpB.

To conclude, the identification of novel RBPs will be a stepping stone for many mechanistic studies. While few studies have tempted to identify and characterize global RNA chaperones in Gram-positive bacteria, using *in vivo* global approaches, post-transcriptional regulation is much less understood than in Gram-negative bacteria. Thus, it is necessary to explore deeper these potential new RBPs and their RNA targets. Characterizing RBPs that regulate RNA-mediated bacterial adaptation, will lead to a better understanding of the physiology and the resilience of *B. subtilis* and Gram-positive in general.

## MATERIAL AND METHODS

### Bacterial strains, plasmids, media and genetic manipulations

All *B. subtilis* strains are derived from wild-type 168 and listed in Table S8. Fragments encompassing *khpA* gene and its own promoter were amplified from genomic DNA, assembled with *Gibson Assembly® Master Mix (NEW ENGLAND BioLabs)* and cloned within pAC5 vector (Martin-Verstraete et al. 1992). A HF tag (HHHHHHRPDYKDDDDK) was added to the C-terminal end of *khpA* by adding the sequence in the reverse primer used for PCR amplification. *khpA* deletion strain was transformed with the pAC5-Khpa-HF plasmid to generate the complementation strain by insertion at the *amyE* locus. Strains were grown either in lysogeny broth (LB) medium or in Difco Sporulation Medium (DSM) supplemented with KCl 0.1%, MgSO4 1 mM, NaOH 0.5 mM, Ca(NO3)2 1 mM, MnCl2 0.01 mM and FeSO4 0.001 mM.

### OOPS (Orthogonal Organic Phase Separation)

To purify the whole RBPome involved in sporulation, we carried out OOPS at defined time points in DSM (3h, 7h, 12h). This approach combines *in vivo* UV crosslinking to stabilize RNA-protein interactions and acidic guanidine phenol chloroform extraction (AGPC, known as TRIzol) to enrich crosslinked protein-RNA complexes. After UV irradiation, the bacterial cells were lysed. The protocol was optimized for *B. subtilis* whose resistant cell wall is mainly composed of peptidoglycan. Then biphasic protein extraction was performed, followed by purification of RBPs. Detailed protocol is are described in supplemental data S1.

### Proteomic sample preparation

All the RBP samples or 1/10^th^ of the free protein samples were denatured with NuPage 1X loading buffer, for 5 min at 65°C and then separated on a NuPage Bolt^TM^ 8% Bis Tris Plus gel for 4 minutes at 150V (the samples migrated as a thin band in the stacking part of the gel). The gels were stained with Coomassie Blue or Imperial Blue for 1 hour, unstained for 3 x 15 min and then overnight before each strip was cut and placed in a low-binding tube. Gel slices were reduced with dithiothreitol, alkylated with iodoacetamide and digested by Trypsin/LysC Mix (Promega) according to the protocol previously described (Belghazi et al. 2024). Samples were analyzed using a hybrid Q-Orbitrap mass spectrometer (Q-Exactive Plus, Thermo Fisher Scientific, USA) coupled to a Vanquish Neo UHPLC system (Thermo Fisher Scientific, USA). Detailed protocol for MS experiment then analysis is described in Supplemental data S1.

### Sporulation efficiency assay

Cultures were inoculated at OD600nm = 0.1 from DSM precultures of each strain and grown in DSM medium for 24 h at 37 °C at 170 rpm. Sporulation efficiency was determined by phase-contrast optical microscopy by counting the number of spores, prespores and vegetative cells from each 24 h DSM cultures in images obtained with a Zeiss Upright Axio Imager M2 microscope. Counts (total >1000) were carried out from 10 to 20 microscopy fields for each experiment. This approach was performed in three biological replicates. Statistically significant differences between strains were determined using a Tukey’s test.

### *In vivo* RNA binding assay

RNA binding assays were performed on strains expressing either KhpA or SpoVR fused to a HF tag, or WT strain for control, grown in LB-medium to transition phase or DSM medium for 7h. UV-crosslinking and lysis were performed as in OOPS. RNA-protein complexes were purified on Magnetic Anti-FLAG M2 beads (Merck). As described in (Delan-Forino and Tollervey 2020), RNAs were partially digested with RNase A/T1 to leave only the “footprint” of the protein. Trimmed complexes were denatured using 6M Guanidinium, immobilized on Ni-NTA affinity resin and washed under denaturing conditions to dissociate copurifying proteins and complexes. Following RNA 5’ labelling with ^32^Phosphate, RNA-protein complexes were eluted and separated on a denaturing SDS-PAGE (NuPAGE Bis-Tris, Invitrogen), transferred on a nitrocellulose membrane and exposed to autoradiography with FLA-5100 Typhoon variable-mode Imager (Amersham Biosciences).

## ACKNOWLEDGEMENTS

We thank the IM2B for financial support (grant for new comers). We thank Regine Lebrun from the Proteomics Platform from the Institut de Microbiologie de la Méditerranée (CNRS, FR3479), which is part of the proteomic network « Marseille Protéomique » (MaP) IBISA and Aix-Marseille Université -labelled, for her expertise. We also thank Jörg Vogel for critical reviewing of the manuscript and Nicolas Waisbord for helpful discussion about correlation analysis.

## Notes

### Competing Interest Statement

The authors have declared no competing interest.

### Summary of Updates

(importance section added, and ref corrected)

## REFERENCES

Abramson J, Adler J, Dunger J, Evans R, Green T, Pritzel A, Ronneberger O, Willmore L, Ballard AJ, Bambrick J, et al. 2024. Accurate structure prediction of biomolecular interactions with AlphaFold 3. Nature 630: 493–500.

Achsel T, Stark H, Lührmann R. 2001. The Sm domain is an ancient RNA-binding motif with oligo(U) specificity. Proc Natl Acad Sci U S A 98: 3685–3689.

Arnaud M, Débarbouillé M, Rapoport G, Saier MH, Reizer J. 1996. In Vitro Reconstitution of Transcriptional Antitermination by the SacT and SacY Proteins of *Bacillus subtilis*. Journal of Biological Chemistry 271: 18966–18972.

Beall B, Moran CP. 1994. Cloning and characterization of spoVR, a gene from *Bacillus subtilis* involved in spore cortex formation. J Bacteriol 176: 2003–2012.

Beckmann BM, Horos R, Fischer B, Castello A, Eichelbaum K, Alleaume A-M, Schwarzl T, Curk T, Foehr S, Huber W, et al. 2015. The RNA-binding proteomes from yeast to man harbour conserved enigmRBPs. Nat Commun 6: 10127.

Belghazi M, Iborra C, Toutendji O, Lasserre M, Debanne D, Goaillard J-M, Marquèze-Pouey B. 2024. High-Resolution Proteomics Unravel a Native Functional Complex of Cav1.3, SK3, and Hyperpolarization-Activated Cyclic Nucleotide-Gated Channels in Midbrain Dopaminergic Neurons. Cells 13: 944.

Birkholz N, Kamata K, Feussner M, Wilkinson ME, Cuba Samaniego C, Migur A, Kimanius D, Ceelen M, Went SC, Usher B, et al. 2024. Phage anti-CRISPR control by an RNA- and DNA-binding helix-turn-helix protein. Nature 631: 670–677.

Boonstra M, Schaffer M, Sousa J, Morawska L, Holsappel S, Hildebrandt P, Sappa PK, Rath H, De Jong A, Lalk M, et al. 2020. Analyses of competent and non-competent subpopulations of *Bacillus subtilis* reveal YhfW, YhxC and ncRNAs as novel players in competence. Environmental Microbiology 22: 2312–2328.

Bresson S, Shchepachev V, Spanos C, Turowski TW, Rappsilber J, Tollervey D. 2020. Stress-Induced Translation Inhibition through Rapid Displacement of Scanning Initiation Factors. Mol Cell 80: 470–484.e8.

Caudron-Herger M, Jansen RE, Wassmer E, Diederichs S. 2021. RBP2GO: a comprehensive pan-species database on RNA-binding proteins, their interactions and functions. Nucleic Acids Research 49: D425–D436.

Chen C, Liu H, Zabad S, Rivera N, Rowin E, Hassan M, Gomez De Jesus SM, Llinás Santos PS, Kravchenko K, Mikhova M, et al. 2021. MoonProt 3.0: an update of the moonlighting proteins database. Nucleic Acids Research 49: D368–D372.

Christopoulou N, Granneman S. 2022. The role of RNA-binding proteins in mediating adaptive responses in Gram-positive bacteria. The FEBS Journal 289: 1746–1764.

Chu L-C, Arede P, Li W, Urdaneta EC, Ivanova I, McKellar SW, Wills JC, Fröhlich T, von Kriegsheim A, Beckmann BM, et al. 2022. The RNA-bound proteome of MRSA reveals post-transcriptional roles for helix-turn-helix DNA-binding and Rossmann-fold proteins. Nat Commun 13: 2883.

Craig JE, Ford MJ, Blaydon DC, Sonenshein AL. 1997. A null mutation in the *Bacillus subtilis* aconitase gene causes a block in Spo0A-phosphate-dependent gene expression. J Bacteriol 179: 7351–7359.

Curtis NJ, Jeffery CJ. 2021. The expanding world of metabolic enzymes moonlighting as RNA binding proteins. Biochemical Society Transactions 49: 1099–1108.

Dai Z, Wu Z, Zhu W, Wu G. 2022. Amino Acids in Microbial Metabolism and Function. In Recent Advances in Animal Nutrition and Metabolism (ed. G. Wu), Vol. 1354 of *Advances in Experimental Medicine and Biology*, pp. 127–143, Springer International Publishing, Cham https://link.springer.com/10.1007/978-3-030-85686-1_7 (Accessed January 22, 2025).

Declerck N, Vincent F, Hoh F, Aymerich S, Van Tilbeurgh H. 1999. RNA recognition by transcriptional antiterminators of the BglG/SacY family: functional and structural comparison of the CAT domain from SacY and LicT. Journal of Molecular Biology 294: 389–402.

Deiorio-Haggar K, Anthony J, Meyer MM. 2013. RNA structures regulating ribosomal protein biosynthesis in bacilli. RNA Biology 10: 1180–1184.

Delan-Forino C, Spanos C, Rappsilber J, Tollervey D. 2020. Substrate specificity of the TRAMP nuclear surveillance complexes. Nat Commun 11: 3122.

Delan-Forino C, Tollervey D. 2020. Mapping Exosome-Substrate Interactions In Vivo by UV Cross-Linking. Methods Mol Biol 2062: 105–126.

D’Halluin A, Gilet L, Lablaine A, Pellegrini O, Serrano M, Tolcan A, Ventroux M, Durand S, Hamon M, Henriques AO, et al. 2025. Embedding a ribonuclease in the spore crust couples gene expression to spore development in *Bacillus subtilis*. Nucleic Acids Research 53: gkae1301.

Dukan S, Nyström T. 1999. Oxidative Stress Defense and Deterioration of Growth-arrested Escherichia coli Cells. Journal of Biological Chemistry 274: 26027–26032.

Durica-Mitic S, Göpel Y, Amman F, Görke B. 2020. Adaptor protein RapZ activates endoribonuclease RNase E by protein-protein interaction to cleave a small regulatory RNA. RNA 26: 1198–1215.

Eichenberger P, Fujita M, Jensen ST, Conlon EM, Rudner DZ, Wang ST, Ferguson C, Haga K, Sato T, Liu JS, et al. 2004. The Program of Gene Transcription for a Single Differentiating Cell Type during Sporulation in *Bacillus subtilis* ed. Jonathan A. Eisen. PLoS Biol 2: e328.

Esteban-Serna S, McCaughan H, Granneman S. 2023. Advantages and limitations of UV cross-linking analysis of protein– RNA interactomes in microbes. Molecular Microbiology 120: 477–489.

Fang S-B, Huang C-J, Huang C-H, Wang K-C, Chang N-W, Pan H-Y, Fang H-W, Huang M-T, Chen C-K. 2017. speG Is Required for Intracellular Replication of *Salmonella* in Various Human Cells and Affects Its Polyamine Metabolism and Global Transcriptomes. Front Microbiol 8: 2245.

Faßhauer P, Busche T, Kalinowski J, Mäder U, Poehlein A, Daniel R, Stülke J. 2021. Functional Redundancy and Specialization of the Conserved Cold Shock Proteins in *Bacillus subtilis*. Microorganisms 9: 1434.

Gamba P, Jonker MJ, Hamoen LW. 2015. A Novel Feedback Loop That Controls Bimodal Expression of Genetic Competence. PLoS Genet 11: e1005047.

González-Gutiérrez JA, Díaz-Jiménez DF, Vargas-Pérez I, Guillén-Solís G, Stülke J, Olmedo-Álvarez G. 2018. The DEAD-Box RNA Helicases of *Bacillus subtilis* as a Model to Evaluate Genetic Compensation Among Duplicate Genes. Front Microbiol 9: 2261.

Goswami B, Nag S, Ray PS. 2024. Fates and functions of RNA-binding proteins under stress. WIREs RNA 15: e1825.

Haldenwang WG, Lang N, Losick R. 1981. A sporulation-induced sigma-like regulatory protein from b. subtilis. Cell 23: 615–624.

Haldenwang WG, Losick R. 1979. A modified RNA polymerase transcribes a cloned gene under sporulation control in *Bacillus subtilis*. Nature 282: 256–260.

Haq IU, Müller P, Brantl S. 2024. A comprehensive study of the interactions in the *B. subtilis* degradosome with special emphasis on the role of the small proteins SR1P and SR7P. Molecular Microbiology 121: 40–52.

Hentze MW, Castello A, Schwarzl T, Preiss T. 2018. A brave new world of RNA-binding proteins. Nat Rev Mol Cell Biol 19: 327–341.

Higgins D, Dworkin J. 2012. Recent progress in *Bacillus subtilis* sporulation. FEMS Microbiol Rev 36: 131–148.

Hilbert DW, Piggot PJ. 2004. Compartmentalization of gene expression during *Bacillus subtilis* spore formation. Microbiol Mol Biol Rev 68: 234–262.

Holmqvist E, Vogel J. 2018. RNA-binding proteins in bacteria. Nat Rev Microbiol 16: 601–615.

Holmqvist E, Wagner EGH. 2017. Impact of bacterial sRNAs in stress responses. Biochem Soc Trans 45: 1203–1212.

Hu LI, Filippova EV, Dang J, Pshenychnyi S, Ruan J, Kiryukhina O, Anderson WF, Kuhn ML, Wolfe AJ. 2018. The spermidine acetyltransferase SpeG regulates transcription of the small RNA rprA ed. F. Rodrigues-Lima. PLoS ONE 13: e0207563.

Hutson M. 2023. Foldseek gives AlphaFold protein database a rapid search tool. Nature d41586-023-02205–4.

Iwańska O, Latoch P, Kopik N, Kovalenko M, Lichocka M, Serwa R, Starosta AL. 2023. Translation in Bacillus subtilis is spatially and temporally coordinated during sporulation. http://biorxiv.org/lookup/doi/10.1101/2023.12.06.569898 (Accessed July 17, 2024).

Jørgensen MG, Pettersen JS, Kallipolitis BH. 2020. sRNA-mediated control in bacteria: An increasing diversity of regulatory mechanisms. Biochim Biophys Acta Gene Regul Mech 1863: 194504.

Khan MA, Durica-Mitic S, Göpel Y, Heermann R, Görke B. 2020. Small RNA-binding protein RapZ mediates cell envelope precursor sensing and signaling in *Escherichia coli*. EMBO J 39: e103848.

Kroos L, Kunkel B, Losick R. 1989. Switch Protein Alters Specificity of RNA Polymerase Containing a Compartment-Specific Sigma Factor. Science 243: 526–529.

Lamm-Schmidt V, Fuchs M, Sulzer J, Gerovac M, Hör J, Dersch P, Vogel J, Faber F. 2021. Grad-seq identifies KhpB as a global RNA-binding protein in *Clostridioides difficile* that regulates toxin production. Microlife 2: uqab004.

Marchais A, Duperrier S, Durand S, Gautheret D, Stragier P. 2011. CsfG, a sporulation-specific, small non-coding RNA highly conserved in endospore formers. RNA Biology 8: 358– 364.

Martin-Verstraete I, Débarbouillé M, Klier A, Rapoport G. 1992. Mutagenesis of the *Bacillus subtilis* “−12, −24” promoter of the levanase operon and evidence for the existence of an upstream activating sequence. Journal of Molecular Biology 226: 85–99.

Matelska D, Steczkiewicz K, Ginalski K. 2017. Comprehensive classification of the PIN domain-like superfamily. Nucleic Acids Research 45: 6995–7020.

Michaux C, Gerovac M, Hansen EE, Barquist L, Vogel J. 2023. Grad-seq analysis of *Enterococcus faecalis* and *Enterococcus faecium* provides a global view of RNA and protein complexes in these two opportunistic pathogens. Microlife 4: uqac027.

Morinaga T, Kobayashi K, Ashida H, Fujita Y, Yoshida K. 2010. Transcriptional regulation of the *Bacillus subtilis* asnH operon and role of the 5′-proximal long sequence triplication in RNA stabilization. Microbiology 156: 1632–1641.

Müller P, Gimpel M, Wildenhain T, Brantl S. 2019. A new role for CsrA: promotion of complex formation between an sRNA and its mRNA target in *Bacillus subtilis*. RNA Biol 16: 972– 987.

Myers TM, Ingle S, Weiss CA, Sondermann H, Lee VT, Bechhofer DH, Winkler WC. 2023. *Bacillus subtilis* NrnB is expressed during sporulation and acts as a unique 3′-5′ exonuclease. Nucleic Acids Research 51: 9804–9820.

Nicolas P, Mäder U, Dervyn E, Rochat T, Leduc A, Pigeonneau N, Bidnenko E, Marchadier E, Hoebeke M, Aymerich S, et al. 2012. Condition-dependent transcriptome reveals high-level regulatory architecture in *Bacillus subtilis*. Science 335: 1103–1106.

Nyström T. 2004. Stationary-Phase Physiology. Annu Rev Microbiol 58: 161–181.

Oda M, Kobayashi N, Fujita M, Miyazaki Y, Sadaie Y, Kurusu Y, Nishikawa S. 2004. Analysis of HutP-dependent transcription antitermination in the *Bacillus subtilis hut* operon: identification of HutP binding sites on *hut* antiterminator RNA and the involvement of the N-terminus of HutP in binding of HutP to the antiterminator RNA. Molecular Microbiology 51: 1155–1168.

Olejniczak M, Jiang X, Basczok MM, Storz G. 2022. KH domain proteins: Another family of bacterial RNA matchmakers? Molecular Microbiology 117: 10–19.

Pandiani F, Chamot S, Brillard J, Carlin F, Nguyen-the C, Broussolle V. 2011. Role of the Five RNA Helicases in the Adaptive Response of *Bacillus cereus* ATCC 14579 Cells to Temperature, pH, and Oxidative Stresses. Appl Environ Microbiol 77: 5604–5609.

Pechter KB, Meyer FM, Serio AW, Stülke J, Sonenshein AL. 2013. Two Roles for Aconitase in the Regulation of Tricarboxylic Acid Branch Gene Expression in *Bacillus subtilis*. J Bacteriol 195: 1525–1537.

Perez-Riverol Y, Bai J, Bandla C, García-Seisdedos D, Hewapathirana S, Kamatchinathan S, Kundu DJ, Prakash A, Frericks-Zipper A, Eisenacher M, et al. 2022. The PRIDE database resources in 2022: a hub for mass spectrometry-based proteomics evidences. Nucleic Acids Research 50: D543–D552.

Pi H, Weiss A, Laut CL, Grunenwald CM, Lin HK, Yi XI, Stauff DL, Skaar EP. 2022. An RNA-binding protein acts as a major post-transcriptional modulator in *Bacillus anthracis*. Nat Commun 13: 1491.

Queiroz RML, Smith T, Villanueva E, Marti-Solano M, Monti M, Pizzinga M, Mirea D-M, Ramakrishna M, Harvey RF, Dezi V, et al. 2019. Comprehensive identification of RNA– protein interactions in any organism using orthogonal organic phase separation (OOPS). Nat Biotechnol 37: 169–178.

Riley EP, Schwarz C, Derman AI, Lopez-Garrido J. 2021. Milestones in *Bacillus subtilis* sporulation research. Microb Cell 8: 1–16.

S. López *, †, Horaci C, Ruzal SM, Sánchez-Rivas C, Rivas EA. 1998. Variations of the Envelope Composition of *Bacillus subtilis* During Growth in Hyperosmotic Medium. Current Microbiology 36: 55–61.

Schujman GE, Grau R, Gramajo HC, Ornella L, De Mendoza D. 1998. *De novo* fatty acid synthesis is required for establishment of cell type-specific gene transcription during sporulation in *Bacillus subtilis*. Molecular Microbiology 29: 1215–1224.

Setlow P, Christie G. 2020. Bacterial Spore mRNA – What’s Up With That? Front Microbiol 11: 596092.

Shchepachev V, Bresson S, Spanos C, Petfalski E, Fischer L, Rappsilber J, Tollervey D. 2019. Defining the RNA interactome by total RNA-associated protein purification. Mol Syst Biol 15: e8689.

Sierro N, Makita Y, De Hoon M, Nakai K. 2008. DBTBS: a database of transcriptional regulation in *Bacillus subtilis* containing upstream intergenic conservation information. Nucleic Acids Research 36: D93–D96.

Smirnov A, Förstner KU, Holmqvist E, Otto A, Günster R, Becher D, Reinhardt R, Vogel J. 2016. Grad-seq guides the discovery of ProQ as a major small RNA-binding protein. Proc Natl Acad Sci USA 113: 11591–11596.

Stragier P. 2022. To Feed or to Stick? Genomic Analysis Offers Clues for the Role of a Molecular Machine in Endospore Formers ed. M.Y. Galperin. J Bacteriol 204: e00187–22.

Tengra FK, Dahl JL, Dutton D, Caberoy NB, Coyne L, Garza AG. 2006. CbgA, a Protein Involved in Cortex Formation and Stress Resistance in *Myxococcus xanthus* Spores. J Bacteriol 188: 8299–8302.

Thomas PD, Ebert D, Muruganujan A, Mushayahama T, Albou L, Mi H. 2022. PANTHER : Making genome-scale phylogenetics accessible to all. Protein Science 31: 8–22.

Trendel J, Schwarzl T, Horos R, Prakash A, Bateman A, Hentze MW, Krijgsveld J. 2019. The Human RNA-Binding Proteome and Its Dynamics during Translational Arrest. Cell 176: 391–403.e19.

Tsimbalyuk S, Shornikov A, Thi Bich Le V, Kuhn ML, Forwood JK. 2020. SpeG polyamine acetyltransferase enzyme from *Bacillus thuringiensis* forms a dodecameric structure and exhibits high catalytic efficiency. Journal of Structural Biology 210: 107506.

Tyanova S, Temu T, Cox J. 2016a. The MaxQuant computational platform for mass spectrometry-based shotgun proteomics. Nat Protoc 11: 2301–2319.

Tyanova S, Temu T, Sinitcyn P, Carlson A, Hein MY, Geiger T, Mann M, Cox J. 2016b. The Perseus computational platform for comprehensive analysis of (prote)omics data. Nat Methods 13: 731–740.

Ul Haq I, Brantl S, Müller P. 2021. A new role for SR1 from *Bacillus subtilis* : regulation of sporulation by inhibition of *kinA* translation. Nucleic Acids Research 49: 10589–10603.

Urdaneta EC, Vieira-Vieira CH, Hick T, Wessels H-H, Figini D, Moschall R, Medenbach J, Ohler U, Granneman S, Selbach M, et al. 2019. Purification of cross-linked RNA-protein complexes by phenol-toluol extraction. Nat Commun 10: 990.

Villanueva E, Smith T, Queiroz RML, Monti M, Pizzinga M, Elzek M, Dezi V, Harvey RF, Ramakrishna M, Willis AE, et al. 2020. Efficient recovery of the RNA-bound proteome and protein-bound transcriptome using phase separation (OOPS). Nat Protoc 15: 2568– 2588.

Vreeland RH, Rosenzweig WD, Powers DW. 2000. Isolation of a 250 million-year-old halotolerant bacterium from a primary salt crystal. Nature 407: 897–900.

Winther AR, Kjos M, Stamsås GA, Håvarstein LS, Straume D. 2019. Prevention of EloR/KhpA heterodimerization by introduction of site-specific amino acid substitutions renders the essential elongasome protein PBP2b redundant in *Streptococcus pneumoniae*. Sci Rep 9: 3681.

Xu T, Li X, Chen K, Qin H, Yi Z, Meng Y, Liu Z. 2022. A Sporulation-Specific sRNA Bvs196 Contributing to the Developing Spore in *Bacillus velezensis*. Microorganisms 10: 1015.

Zhang Y, Burkhardt DH, Rouskin S, Li G-W, Weissman JS, Gross CA. 2018. A Stress Response that Monitors and Regulates mRNA Structure Is Central to Cold Shock Adaptation. Molecular Cell 70: 274–286.e7.

Zheng JJ, Perez AJ, Tsui HT, Massidda O, Winkler ME. 2017. Absence of the KhpA and KhpB (JAG/EloR) RNA-binding proteins suppresses the requirement for PBP2b by overproduction of FtsA in *Streptococcus pneumoniae* D39. Molecular Microbiology 106: 793–814.

Zhou B, Xiong Y, Nevo Y, Kahan T, Yakovian O, Alon S, Bhattacharya S, Rosenshine I, Sinai L, Ben-Yehuda S. 2023. Dormant bacterial spores encrypt a long-lasting transcriptional program to be executed during revival. Molecular Cell 83: 4158–4173.e7.

Zhu B, Stülke J. 2018. SubtiWiki in 2018: from genes and proteins to functional network annotation of the model organism *Bacillus subtilis*. Nucleic Acids Res 46: D743–D748.

Zhu Y, Ponath F, Cosi V, Vogel J. 2024. A global survey of small RNA interactors identifies KhpA and KhpB as major RNA-binding proteins in *Fusobacterium nucleatum*. Nucleic Acids Res 52: 3950–3970.

Zuberi AR, Doi RH. 1990. A mutation in P23, the first gene in the RNA polymerase sigma A (sigma 43) operon, affects sporulation in *Bacillus subtilis*. J Bacteriol 172: 2175–2177.

