## Supplementary material for "Remodeling of RNA-Binding Proteome and RNA-mediated regulation as a new layer of control of sporulation": Sup Figures S1, S2, S3

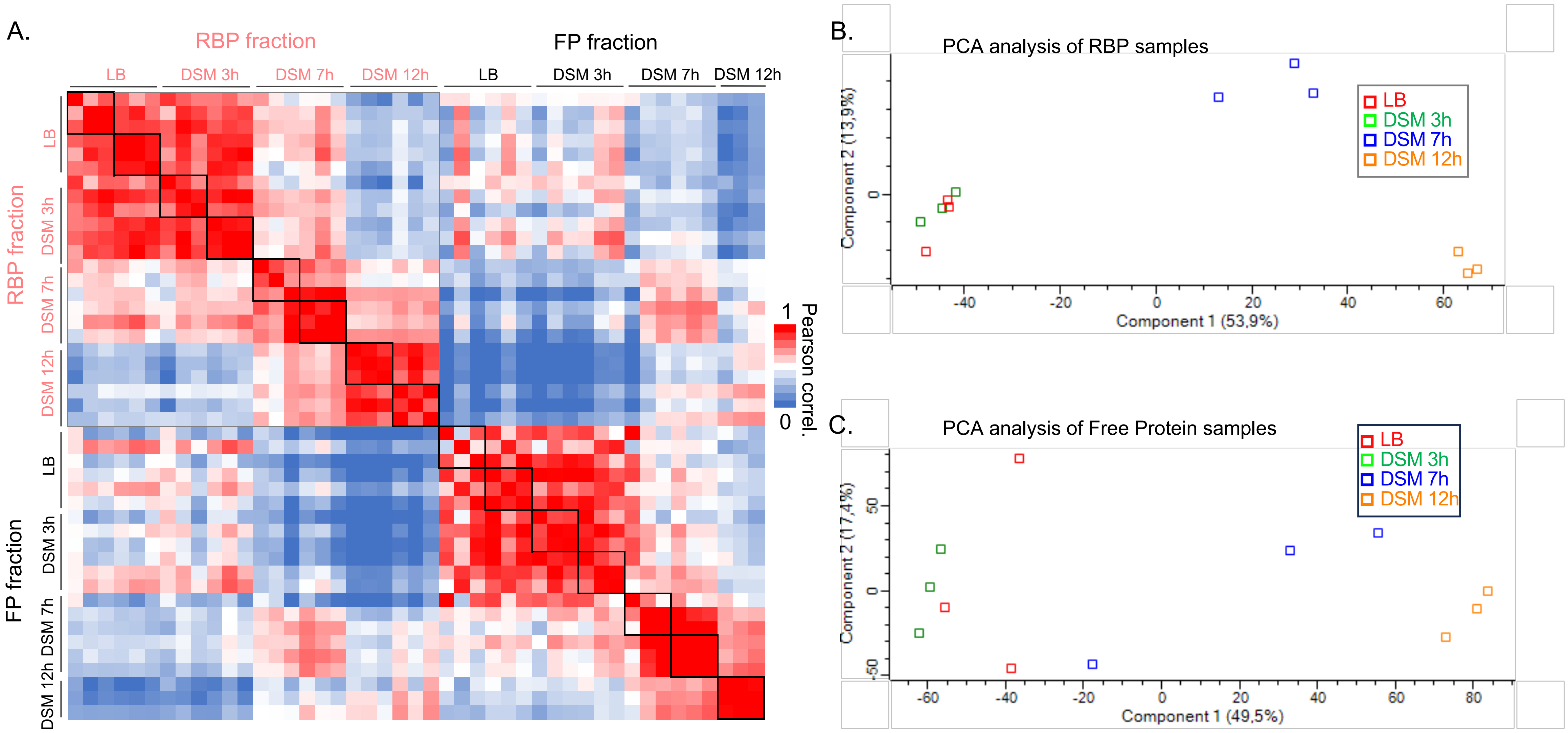


Kaboré *et al*, Sup Figure S1

Fig. S1: A. Correlation matrix displayed as a heat map showing Pearson correlation between datasets (with LFQ intensities) obtained for the individual replicates of RBP and free protein (FP) fractions, in LB and along sporulation process in DSM medium. B-C. Principal component analysis of triplicates for each condition for RBP (B) and free proteins fractions (C) samples.

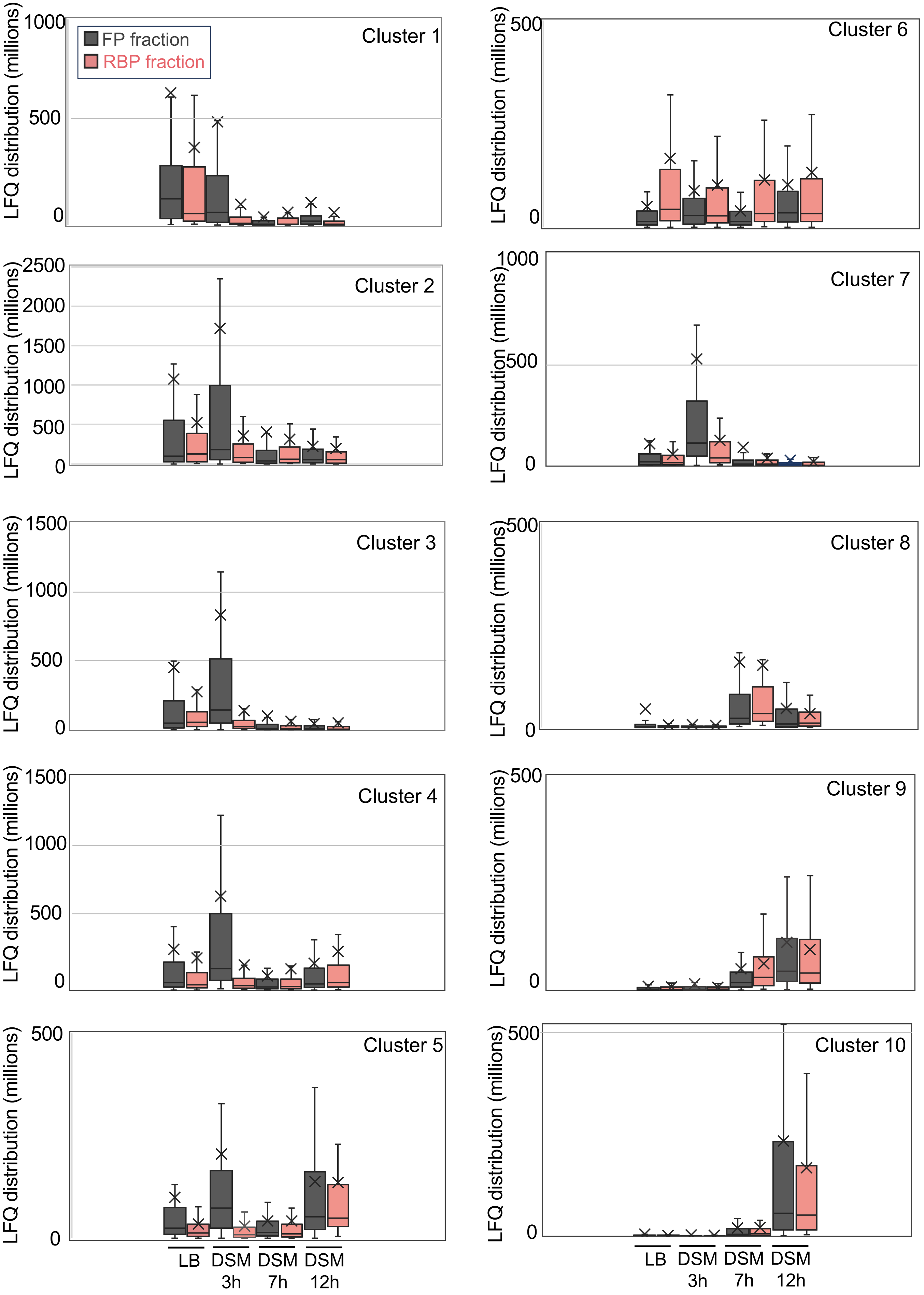


Kaboré *et al*, Sup Figure S3

Fig. S3: Box plots exhibiting distributions of LFQ intensities of proteins within each cluster (from Fig. 7) in both free protein and RBP fractions.

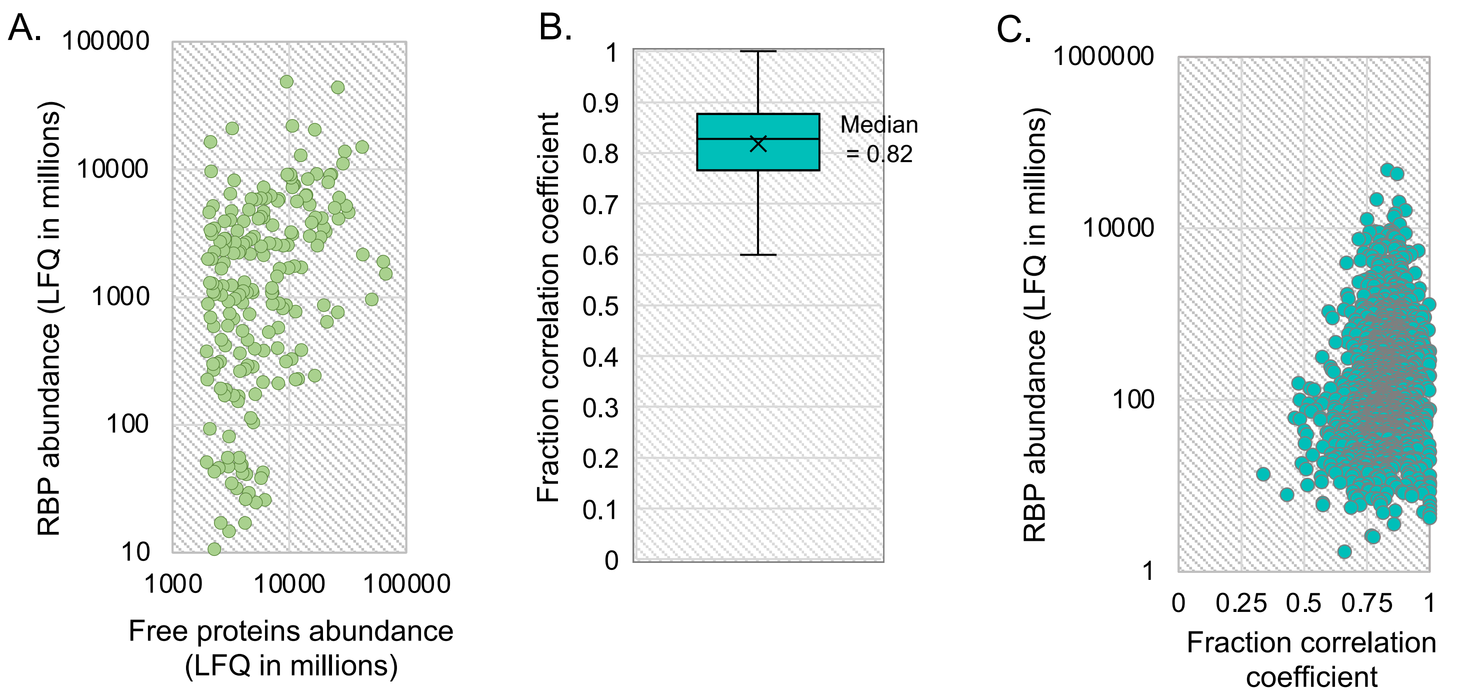


Kaboré *et al*, Sup Figure S2

Fig. S2: A. LFQ intensities averaged for both RBP and free protein fractions for the top 100 proteins identified in the free protein fraction were displayed on a 2D-Scatter plot. B. Distribution of fraction correlation coefficients calculated for all 2102 proteins (free proteins and RBPs). C. LFQ scores averaged from RBP fractions and dynamic correlation coefficients of all 2102 proteins were displayed on a 2D-Scatter plot.
