## Supplementary material for "Remodeling of RNA-Binding Proteome and RNA-mediated regulation as a new layer of control of sporulation": Sup MM + Sup figures legends

**Supplemental data S1**

**SUPPLEMENTAL MATERIAL & METHODS**

**OOPS (Orthogonal Organic Phase Separation)**

*UV crosslink*: 200 ml cultures in DSM medium were inoculated at $OD_{600nm}$=0.1 from DSM pre-cultures at $OD_{600nm}$=1. $OD_{600nm}$ was measured at 3 h, 7 h, 12 h. A volume of culture equivalent to 200 OD units was filtered through a cellulose membrane that was then placed on a metal block cooled to -70°C and cut into 2 equal parts. One half was exposed to UV (λ = 254 nm; 1x9999x 100 µJ/cm2) (UV+) to induce covalent bonds between proteins and RNAs in direct interaction. The other half of the sample serves as a control not subjected to UV (UV-). The bacteria on the membranes were then resuspended in PBS (Phosphate-Buffered Saline) buffer and centrifuged for 5 min at 1800 RCF at 4°C. The pellets were stored at -80°C.

*Lysis:* bacterial pellets were then resuspended in 50 µl of TN150 buffer (50 mM Tris, pH 7.5, 150 mM NaCl), supplemented with 500 µg lysozyme (to degrade peptidoglycan) and 12 units of Promega DNase RQ1 (to degrade DNA and dissociate protein-DNA complexes). After incubation for 30 minutes at 37°C, 150 µl of Trizol and 550 mg of beads (lysis matrix B, MP Biomedical) were added and bacteria were mechanically lysed (Fastprep, MP Biomedical 6 x 45s at 6.5 m/s). After lysis, 1 ml of TRIzol was added to the samples. Samples were then centrifuged for 5 min at 1800 RCF at 4°C and supernatants transferred to new tubes for storage at -20°C.

*Biphasic extraction*: after the addition of chloroform (1:5), the samples were vortexed and centrifuged for 15 min at 12,000 RCF to form three distinct phases: the aqueous phase containing the free RNAs, the interphase containing the RBP-RNA complexes and the organic phase containing the free proteins. These 3 phases were collected separately. The free RNA and free protein phases were stored at -20°C. The interphase was subjected to a second phase separation in which the aqueous and organic phases were eliminated, leaving only the interphase. This prevents free protein contamination in the remaining organic phase.

Interphases were washed successively with methanol (9:1), centrifuged for 10 min at 23,000 RCF at 4°C and stored at -20°C.

*To purify RBPs in complex with RNAs*, pellets were resuspended in 100 µl of 100 mM TEAB (trimethylammonium bicarbonate) containing 1% SDS. To degrade completely the RNAs, the samples were sonicated for 15 min (high intensity, 30 s pause/cycle), incubated for 20 min at 95°C with 1 mM $MgCl_2$ to optimize hydrolysis, then incubated in the presence of 1 µl RNase A + T1 (RNAce-IT Ribonuclease Cocktail, Agilent) at 37°C for 3h. This step was repeated once more with a 16 h incubation. After degradation of the RNAs, a phase separation was carried out as before. In this step, the organic phase containing the RBPs previously bound to the RNAs was transferred to a low-binding tube. Proteins were precipitated in the presence of pure ethanol, centrifuged for 10 min at 23,000 RCF at 4°C, then washed with 80% ethanol and centrifuged for 10 min at 23,000 RCF at 4°C and the pellets were resuspended in 20 µl of ultra-pure $H_20$. The phase containing the free proteins previously stored at 20°C during the biphasic extraction above is processed similarly.

*To purify proteins bound RNAs (PBR),* part of the interphase (1/3) is kept. Samples were treated with proteinase K solution (1.3 mg/mL, enzyme, Tris HCl, pH8 and 10mM EDTA) for 2 h at 50°C to degrade RNA-associated proteins. Next, 300 µl of phenol:chloroform:isoamyl alcohol mixture (vol/vol, 25:24:1) was added to dissociate the RNAs from the degraded proteins. After centrifugation for 15 min at 12,000 RCF at 4°C, the aqueous phase containing the RNAs was transferred to a new tube.

RNAs (free and PBR) were precipitated for 10 min on ice in the presence of 1:10 volume of sodium acetate (3M, pH 5.3) and 1:2 volumes of 100% isopropanol. After centrifugation at 23,000 RCF for 10 min at 4°C, the pellet was washed with 900 µl of 100% ethanol followed by 900 µl of 70% ethanol. After removing the supernatant and allowing the remaining ethanol to evaporate, the pellets were resuspended in ultrapure water. RNA purity and concentrations were determined using Nanodrop (Thermo Fisher Scientific). To visualize the integrity of the purified RNAs, samples were placed on a 1% agarose gel after being denatured for 10 min at 65°C (95% formamide 150 mM EDTA 0.025% xylene cyanol and bromophenol).

**Mass spectrometry**

Peptides were trapped on a C18 PepMap 300 trap column (300 μm × 5 mm, 5 μm, 300 Å) and separated on an EASY-Spray™ PepMap™ Neo capillary column (75 μm × 50 cm or 75 μm × 15 cm, 2 μm, 100 Å) at a flow rate of 250 nL/min. The mobile phases consisted of 0.1% formic acid in water (A) and 80% acetonitrile/20% water with 0.1% formic acid (B). The gradient elution was from 2% to 25% B over 90 minutes, followed by an increase to 50% B over 20 minutes. MS spectra were acquired at a resolution of 70,000 over a mass range of 350–1,900 m/z, with a maximum injection time of 100 ms. Fragmentation spectra of the 10 most abundant precursors with charge states ≥2 (Top10 method in Data-Dependent Acquisition mode) were recorded using high-energy collision dissociation (HCD) with a normalized collision energy of 27%. Raw data were processed and searched against the *B. subtilis* database (Taxonomy ID: 224308, downloaded from UniProt on 2022-10-17) using Proteome Discoverer 3.0 (Thermo Fisher Scientific, USA). Protein identification was performed with the Sequest HT algorithm using the following parameters: dynamic modifications—methionine oxidation, N-terminal acetylation, and N-terminal methionine loss; static modification—carbamidomethylation of cysteines; mass tolerances—10 ppm for precursor ions and 0.02 Da for fragment ions; and up to three missed tryptic cleavages. Label-free quantification was based on precursor ion intensities, with normalization to the total peptide amount.

**Proteomics analysis**

MS/MS data were processed using MaxQuant (v.2.5.0.0; downloaded from https://maxquant.net/maxquant/, Cox lab, Max Planck institute of Biochemistry, Martinsried, Germany) for Label Free Quantification (Tyanova et al. 2016a). The databank was the same as the one used for Proteome Discoverer analysis. Maximum missed cleavages by trypsin were set to 3, carbamidomethyl of Cysteines was set as fixed modification and oxidation of Methionines and acetylation of N-terminal were variable.  Match between runs was used with a match time window of 0.7 min and an alignment time window of 20 minutes. Otherwise, all other parameters were left to default.

The ProteinGroup.txt file generated by Maxquant was uploaded in Perseus software (v 1.6.15) (Tyanova et al. 2016b) downloaded from https://maxquant.net/perseus. We applied quality filters to preserve only the proteins robustly identified in at least three samples from one of the replicates (See Fig. 3C). Missing values were replaced from normal distribution. The proteins showing a ratio CL+/CL- lower than 1 and proteins annotated as secreted and glycoproteins could be found in the interphase non-specifically and were filtered out. A fraction correlation coefficient was calculated for each protein recovered in OOPS to assess the differences between recovery of one protein on both fractions, according to this formula: Fraction correlation coefficient = $1 - (\sqrt{(0.25 \times (FP_{LB} - RBP_{LB})^2 + 0.25 \times (FP_{DSM\ 3h} - RBP_{DSM\ 3h})^2 + 0.25 \times (FP_{DSM\ 7h} - RBP_{DSM\ 7h})^2 + 0.25 \times (FP_{DSM\ 12h} - RBP_{DSM\ 12h})^2})$. The mass spectrometry proteomics data have been deposited to the ProteomeXchange consortium via the Pride Proteomics Identification Database partner repository with the data set identifier PXD061929 (Perez-Riverol et al. 2022).

### Supplementary Figures legends

Fig. S1: A. Correlation matrix displayed as a heat map showing Pearson correlation between datasets (with LFQ intensities) obtained for the individual replicates of RBP and free protein (FP) fractions, in LB and along sporulation process in DSM medium. B-C. Principal component analysis of triplicates for each condition for RBP (B) and free protein (C) samples.

Fig. S2: A. LFQ intensities averaged for both RBP and free protein fractions for the top 100 proteins identified in the free protein fraction were displayed on a 2D-Scatter plot. B. Distribution of fraction correlation coefficients calculated for all 2102 proteins (free proteins and RBPs). C. LFQ scores averaged from RBP fractions and dynamic correlation coefficients of all 2102 proteins were displayed on a 2D-Scatter plot.

Fig. S3: Fig. S3: Box plots exhibiting distributions of LFQ intensities of proteins within each cluster (from Fig. 7) in both free protein (FP) and RBP fractions.

**Sup Tables:**

Table S1: Normalized quantities of putative sRNAs recovered by microarray in vegetative growth (LB) and during sporulation, used for Fig. 2. Data from Nicolas et al, 2012.

Table S2: Matrix exhibiting all Pearson correlation coefficients between OOPS replicates. Correlation was calculated on datasets composed of 2102 proteins (without replacing missing values). It was used for Fig. 4C and D and Fig. S1A).

Table S3: The 2102 proteins recovered in OOPS (in at least one replicate) were sorted by their LFQ intensities in the free protein fraction (averaged from 3 individual biological replicates) in column 2. Corresponding LFQ intensities from RBP fraction and fraction correlation coefficient are shown in columns 3 and 4 respectively. This table was used for Fig. 5A and B.

Table S4: List of proteins found by OOPS in RBP fraction in each condition (LB, DSM 3 h, 7 h, 12 h). Only proteins being present in at least the 3 individual replicates of RBP fractions from a same condition, not known as being glycosylated or secreted, and exhibiting a ratio UV+/UV- superior at 1 were kept (1520 proteins in total). This list was used for Venn diagram in Fig. 6.

Table S5:

Proteins recovered in OOPS, normalized LFQ were clustered by Euclidian distance used for Fig. 5B. Enriched GO-terms were identified using Panther Gene Ontology database (Thomas et al. 2022); with false discovery rate (FDR)-adjusted p-value lower than 0.05 except when stated otherwise in cluster 10.

Table S6:

RNases and effectors of RNA degradation recovered in OOPS

Table S7:

Proteins recovered in OOPS and found in clusters 7 and 8.

Table S8:

List of strains used in this study.

Table S9:

RBP abundance (LFQ scores) recovered in OOPS.

Table S10:

List of RBPs containing HTH and Sm/LSm domains recovered in OOPS.
